# Combining environmental DNA and remote sensing for efficient, fine-scale mapping of arthropod biodiversity

**DOI:** 10.1101/2023.09.07.556488

**Authors:** Yuanheng Li, Christian Devenish, Marie I. Tosa, Mingjie Luo, David M. Bell, Damon B. Lesmeister, Paul Greenfield, Maximilian Pichler, Taal Levi, Douglas W. Yu

**Affiliations:** State Key Laboratory of Genetic Resources and Evolution and Yunnan Key Laboratory of Biodiversity and Ecological Security of Gaoligong Mountain, Kunming Institute of Zoology, Chinese Academy of Sciences, Kunming, Yunnan, China 650223; Faculty of Biology, University of Duisburg-Essen, Essen, Germany D-45141; School of Biological Sciences, University of East Anglia, Norwich Research Park, Norwich, Norfolk, UK NR47TJ; Department of Fisheries, Wildlife, and Conservation Sciences, Oregon State University, Corvallis, Oregon USA 97331; Kunming College of Life Sciences, University of Chinese Academy of Sciences, Kunming, China; Pacific Northwest Research Station, U.S. Department of Agriculture Forest Service, Corvallis, OR, USA 97331; CSIRO Energy, Lindfield, NSW, Australia; School of Biological Sciences, Macquarie University, Australia; Theoretical Ecology, University of Regensburg, Regensburg, Germany; Center for Excellence in Animal Evolution and Genetics, Chinese Academy of Sciences, Kunming Yunnan, China 650223; School of Geography, Geology and the Environment, Keele University, Staffordshire, ST5 5BG, UK

**Keywords:** metabarcoding, environmental DNA, metagenomics, Earth Observation, biodiversity indices, Arthropoda, site irreplaceability, systematic conservation planning, conservation, forestry, machine learning, joint species distribution model

## Abstract

Arthropods contribute importantly to ecosystem functioning but remain understudied. This undermines the validity of conservation decisions. Modern methods are now making arthropods easier to study, since arthropods can be mass-trapped, mass-identified, and semi-mass-quantified into ‘many-row (observation), many-column (species)’ datasets, with homogeneous error, high resolution, and copious environmental-covariate information. These ‘novel community datasets’ let us efficiently generate information on arthropod species distributions, conservation values, uncertainty, and the magnitude and direction of human impacts. We use a DNA-based method (barcode mapping) to produce an arthropod-community dataset from 121 Malaise-trap samples, and combine it with 29 remote-imagery layers within a joint species distribution model. With this approach, we generate distribution maps for 76 arthropod species across a 225 km^2^ temperate-zone forested landscape. We combine the maps to visualise the fine-scale spatial distributions of species richness, community composition, and site irreplaceability. Old-growth forests show distinct community composition and higher species richness, and stream courses have the highest site-irreplaceability values. With this ‘sideways biodiversity modelling’ method, we demonstrate the feasibility of biodiversity mapping at sufficient spatial resolution to inform local management choices, while also being efficient enough to scale up to thousands of square kilometres.

## INTRODUCTION

Arthropods contribute in numerous ways to ecosystem functioning (Prather et al., 2013) but are understudied relative to vertebrates and plants (Troudet et al., 2017). This taxonomic bias undermines the validity of conservation decisions when the effects of change in climate, land use, and land cover differ across taxa (Hamilton et al., 2022; Westgate et al., 2014). Also, it is arguable that modern methods now make arthropods *easier* to study than vertebrates and plants, given that arthropods can be mass-trapped and mass-identified (Chua et al., 2023; van Klink et al., 2022). Another logistical advantage is that arthropod community structure is correlated with vegetation structure (Lewinsohn and Roslin, 2008; Zhang et al., 2016), and since vegetation can be measured remotely at large spatial scale via airborne and spaceborne sensors (Bush et al., 2017), remote imagery could also provide large-spatial-scale information on arthropods. In fact, it is already known that spaceborne SAR (synthetic aperture radar) and airborne lidar (Light Detection And Ranging) imagery of fine-scale forest structure can predict the distributions of entomofauna and avifauna (Bae et al., 2019; Müller et al., 2009; Müller and Brandl, 2009; Rhodes et al., 2022).

### Successful governance of the biodiversity commons

Arthropod conservation should be seen in the wider context of efficient biodiversity governance. Dietz et al.’s (2003) framework for the successful governance of public goods can be usefully summarised into five elements: (1) information generation, (2) infrastructure provision, (3) political bargaining, (4) enforcement, and (5) institutional redesign. The first element, information generation, asks engineers and scientists to generate *high-quality, granular, timely, trustworthy*, and *understandable* information on ecosystem status and change, values, uncertainty, and the magnitude and direction of human impacts.

Although there exists an example of the five elements working together to achieve single-species conservation (see Supplementary Information: “Dietz et al.’s five elements”), to our knowledge, there is so far no example of the five elements comprehensively working together to achieve *multi-species* conservation, in large part because the tools, study designs, and analyses needed to generate information on many species at once are complex. This complexity is a barrier to uptake, delaying the institutional redesigns that could operationalise, finance, and scale-up conservation.

Our focus in this study is therefore to demonstrate how to efficiently generate *high-quality, granular, timely, trustworthy*, and *understandable* information on status and change in arthropod biodiversity, conservation value, uncertainty, and the magnitude and direction of human impacts.

We use the management of National Forests in the United States as our test case for multi-species biodiversity conservation. This management should follow the doctrine outlined in the 1960 Multiple-Use Sustained-Yield Act that requires management and utilisation of natural resources to satisfy multiple competing interests and to maintain the natural resources in perpetuity (Carter et al., 2019; Hobbs et al., 2010; Loomis, 2002). Although US law mandates that each use be given equal priority, implementation is stymied by a lack of biodiversity data such as distribution maps of large numbers of species to identify areas of high conservation value that can be protected while still supporting extractive uses in other areas. Moreover, the species distribution maps should be regularly updated so that the impacts of management interventions can be inferred, feeding back to adaptive management (Frankham, 2010; Bush et al., 2017).

### High-throughput arthropod inventories

Now though, there are new technologies capable of efficiently and granularly capturing biodiversity information, via DNA isolated from environmental samples (eDNA) and via electronic sensors (bioacoustics, cameras, radar) (Besson et al., 2022; Bohmann et al., 2014; Bush et al., 2017; Christin et al., 2019; Pawlowski et al., 2020; Ruppert et al., 2019; Tosa et al., 2021; van Klink et al., 2022; Chua et al., 2023). The eDNA methods start with DNA-based taxonomic assignment (‘DNA barcoding’ Hebert et al., 2003) and vary in how the DNA is collected and processed. For instance, large numbers of arthropods can efficiently be individually DNA-extracted and sequenced to produce count datasets (Ratnasingham, 2019; Srivathsan et al., 2021). These DNA-barcoded specimens (plus human-identified specimens) can optionally be used to annotate specimen images to train deep-learning models to scale up identifications (Chua et al., 2023; van Klink et al., 2022). Alternatively, DNA from arthropods can be extracted *en masse* from traps (Ji et al., 2013) or from environmental substrates, such as water washes of flowers (e.g. Thomsen and Sigsgaard, 2019) and mass-sequenced. These latter processing pipelines are known as ‘metabarcoding’ or ‘metagenomics’, depending on whether the target DNA-barcode sequence is PCR-amplified (both described in Bush et al., 2017).

The eDNA- and sensor-based methods can all produce ‘novel community data’, which Hartig et al. (2023) describe as ‘many-row (observation), many-column (species)’ datasets, therefore making possible high spatial and/or temporal resolution and extent. Novel community data contain some form of abundance information, ranging from counts to within-species abundance change (Luo et al., 2023; Diana et al., 2022) to presence/absence, and because the methods are automated and standardised, the errors in these datasets tend to be homogeneous (e.g. minimal observer effects), which facilitates their correction given appropriate sample replicates and statistical models.

### ‘Sideways’ biodiversity modeling and site irreplaceability ranking

It is natural to think about combining novel community data with copious environmental-covariate information in the form of continuous-space remote-imagery layers (and/or with continuous-time acoustic series) to produce continuous spatio(-temporal) biodiversity data products (Bush et al., 2017; He et al., 2015; Kwok, 2018; Leitão and Santos, 2019; Lin et al., 2021; Pettorelli et al., 2018; Cavender-Bares et al., 2022; Hartig et al., 2023; Müller et al., 2023; Davis et al., 2023). Here we do just this, combining a point-sample dataset of Malaise-trapped arthropods with continuous-space Landsat and lidar imagery within a joint species distribution model (JSDM Ovaskainen and Abrego, 2020; Pichler and Hartig, 2021; Warton et al., 2015; Davis et al., 2023). We were able to produce distribution maps for 76 arthropod species across a forested landscape. Because this landscape is characterised by overlapping gradients of environmental conditions (e.g. elevation, distance from streams and roads) and mosaics of management (e.g. clearcuts, old-growth), we can estimate the effects of different combinations of natural and anthropogenic drivers on arthropod biodiversity, including combinations that were not included in our sample set. We can also subdivide the landscape into management units and rank them by conservation value, to inform decision-making in this multi-use landscape.

The above approach is a direct test of a protocol originally proposed by Bush et al. (2017) and more formally described by Pollock et al. (2020) under the name ‘sideways’ biodiversity modelling. In short, sideways biodiversity models (1) integrate “the largely independent fields of biodiversity modeling and conservation” and (2) include large numbers of species in conservation planning instead of using habitat-based metrics. Or in plain language, we use remote-sensing imagery to fill in the blanks between our sampling points, which creates a continuous map of arthropod biodiversity that we can use to study arthropod ecology and guide conservation.

## MATERIALS AND METHODS

In short, we combine DNA-based species detections, remote-sensing-derived environmental predictors, and joint species distribution modelling to predict and visualise the fine-scale distribution of arthropods across a large forested landscape. We use the joint predictions from the JSDM to map species richness, compositional distinctiveness, and conservation value across the landscape. For the detailed protocol and explanations of the field, laboratory, bioinformatic, and statistical methods, see Supplementary Information: Materials and Methods.

### Model Inputs

#### Field data collection

We collected with 121 Malaise-trap samples for seven days into 100% ethanol at 89 sampling points in and around the H.J. Andrews Experimental Forest (HJA), Oregon, USA in July 2018 (Figure 1). Sites were stratified by elevation, time since disturbance, and inside and outside the HJA (inside = a long-term research site with no logging since 1989; outside = continued active management). HJA represents a range of previously logged to primary forest, but with notably larger areas of mature and old-growth forest reserves than the regional forest mosaic, which consists of short-rotation plantation forests on private land and a recent history of active management on public land.

**Figure 1.**
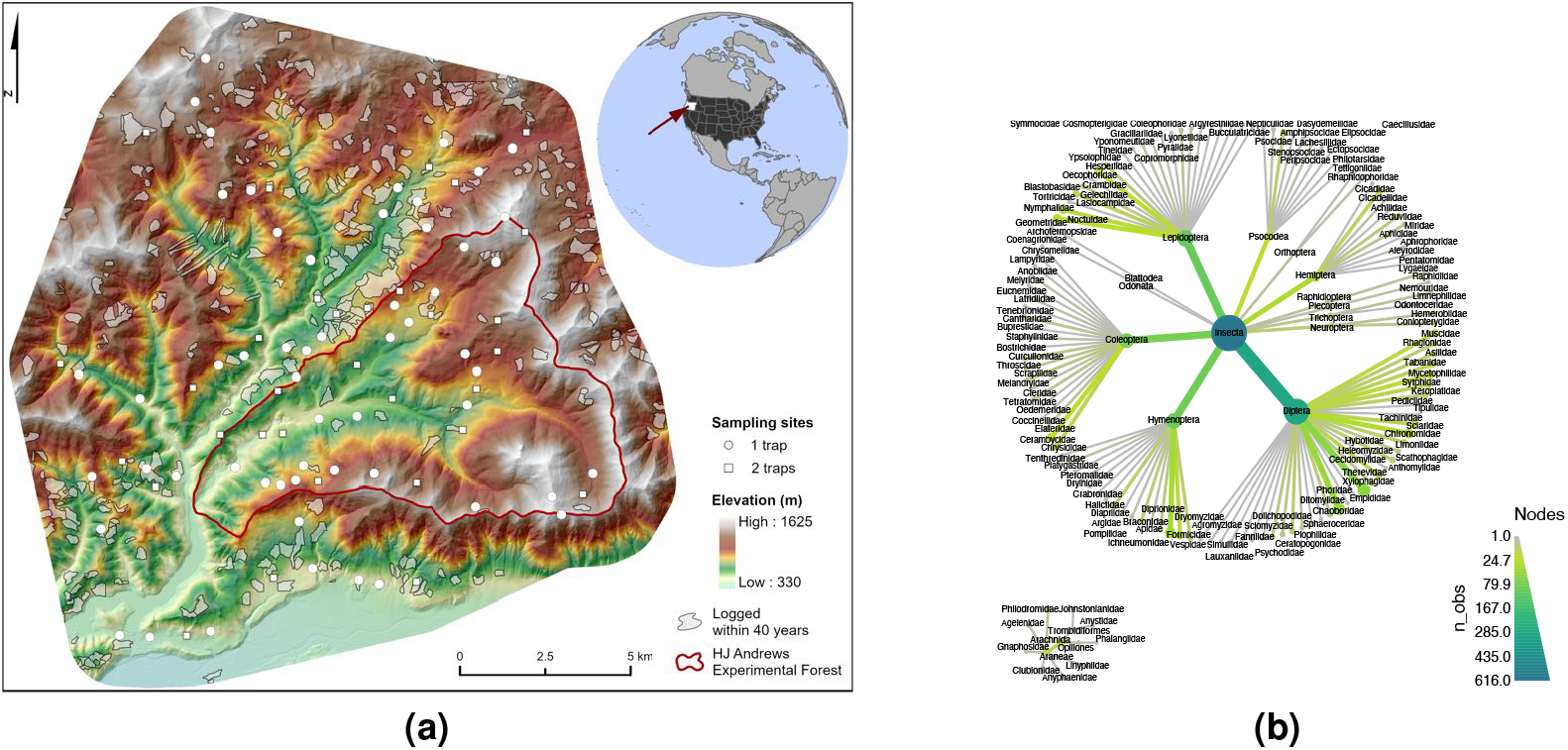
Sampling design and taxonomic diversity of the Malaise trapping campaign. (a). Sampling points in and around the H.J. Andrews Experimental Forest (red line), Oregon, USA. The study area consists of old-growth and logged (gray patches) deciduous and evergreen forest under different management regimes. Arthropods were sampled with Malaise traps at 89 sampling points in July 2018, with one trap at 57 points (white circles) and with two traps 40 m apart at 32 points (white squares). Elevation indicated with a green to white false-color gradient. (b). Taxonomic distribution of all detected Operational Taxonomic Units (OTUs) from the samples. Node size and color are scaled to the number of OTUs. See Figure 4S for a heat tree of the 190 included OTUs.

#### Wet-lab pipeline and bioinformatics

##### DNA extraction and sequencing

We extracted the DNA from each Malaise-trap sample by soaking the arthropods in a lysis buffer and sent it to Novogene (Beijing, China) for whole-genome shotgun sequencing.

#### Creating a barcode reference database using Kelpie in-silico PCR

On the output fastq files, we carried out ‘in-silico’ PCR using `Kelpie 2.0.11` Greenfield et al. (2019) and the BF3+BR2 primers from Elbrecht et al. (2019), outputting 5560 unique DNA-barcode sequences.

After 97%-similarity clustering and filtering for erroneous sequences, we were left with 1225 OTUs as the reference barcode set.

#### Read mapping to reference barcodes

We then mapped the reads of each sample to the reference barcodes, creating a 121-sample *×* 1225-OTU table. A species was accepted as being in a sample if reads mapped at high quality along more than 50% of its barcode length, following acceptance criteria from Ji et al. (2020).

#### Environmental covariates

To predict species occurrences in the areas between the sampling points, we collected 58 continuous-space predictors (Table 1S), relating to forest structure, vegetation reflectance and phenology, topography, and anthropogenic features, restricting ourselves to predictors that can be measured remotely. The forest-structure variables were from airborne lidar data collected from 2008 to 2016, which correlate with forest structure in US Pacific Northwest coniferous forests, such as mean diameter, canopy cover, and tree density (Kane et al., 2010). The vegetation-related variables came from Landsat 8 individual bands, plus standard deviation, median, 5% and 95% percentiles of those bands over the year, and indices of vegetation status e.g. Normalized Difference Vegetation Index (NDVI). Both the proportion of canopy cover and annual Landsat metrics were calculated within radii of 100, 250 and 500 m, given that vegetation structure at different spatial scales is known to drive arthropod biodiversity (Müller et al., 2014). The topography variables were calculated from lidar ground returns, including elevation, slope, Eastness and Northness split from aspect, Topographic Position Index (TPI), Topographic Roughness Index (TRI) (Wilson et al., 2007), Topographic Wetness Index (TWI) (Metcalfe et al., 2018), and distance to streams, based on a vector stream network (http://oregonexplorer.info, accessed 24 Oct 2019). The anthropogenic variables include distance to nearest road, proportion of area logged within the last 100 and within the last 40 years, within radii of 250, 500, and 1000 m, and a categorical variable of inside or outside the boundary of the H.J. Andrews Experimental Forest. They are not directly derived from remote-sensing data, but we included them because they could be derived from remote-sensing imagery. We then reduced our 58 environmental covariates to 29, removing the covariates that were most correlated with the others (as measured by Variance Inflation Factor). The 29 retained covariates include six anthropogenic activities, two raw Landsat bands, seven indices based on annual Landsat data, six canopy/vegetation-related variables from LiDAR, and eight topography variables (Table 1S, Figure 5S), which we mapped across the study area at 30 m resolution.

### Statistical Analyses

#### Species inputs

We converted the sample *×* species table to presence-absence data (1*/*0), and we only included species present at ≥ 6 sampling sites across the 121 samples. Our species dataset was thus reduced to 190 species in two classes, Insecta and Arachnida (Figure 1b).

#### Joint Species Distribution Model

The general idea behind species distribution modelling is to ‘predict a species’ distribution’. We use each species’ observed incidences (1*/*0) at all sampling points, plus the environmental-covariate values at those points, to ‘fit’ a model that predicts the species’ incidences from the covariate values. Once we have a fitted model, we use it to predict the species’ probability of presence over the rest of the sampling area, where the environmental-covariate values are known but the species’ incidences are not. Spatial autocorrelation was accounted by a trend-surface component. Joint species distribution models (JSDM) extend individual SDMs by additionally accounting for co-occurrences of species (See Supplementary Information: Joint Species Distribution Model).

#### Tuning and testing

The statistical challenge is to avoid overfitting, which is when the fitted model does a good job of predicting the species’ incidences at the sampling points that were used to fit the model in the first place but does a bad job of predicting the species over the rest of the landscape. Overfitting is likely in our dataset because many of our species are rare, there are many candidate remote-sensing covariates, and we expect that any relationships between remote-sensing-derived covariates and arthropod incidences are indirect and thus complex, necessitating the use of flexible mathematical functions.

To minimise overfitting, we used regularisation and cross-validation. Regularisation uses penalty terms during model fitting to favour a relatively simple set of covariates, and cross-validation finds the best values for those penalty terms (tuning). First, we randomly split the species incidence data from the 121 samples in 89 sampling points into 75% training data (*n* = 91) and 25% test data (*n* = 30) (Figure 1S). The training data was used to try 1000 different hyperparameter combinations in a five-fold cross-validation design, some of which are the penalty terms, to find the combination that achieves the highest predictive performance on the training data itself (see Supplementary Information: Tuning and Testing, Figure 1S). The model with this combination was then applied to the 25% test data to measure true predictive performance. To fit the model, we used the JSDM `R` package `sjSDM 1.0.5` (Pichler and Hartig, 2021), with the DNN (deep neural network) option to account for complex, nonlinear effects of environmental covariates (the DNN outperformed a linear model, see Figure 11S), which suits our dataset of many species with few data points and many covariates.

Finally, to estimate how OTU incidence affects the variability of predictive accuracies, we also tuned a model to the whole dataset in a five-fold cross-validation, found optimal hyperparameters, and used them in another five-fold cross-validation on the entire dataset to estimate the variability of predictive AUCs by OTU (see Supplementary Information: Variability in Predictive AUC by OTU Incidence). We emphasise that method is only useful for estimating variability in predictive performance, given that it potentially overestimates predictive performance, which is what we avoided by using a pure holdout in the main analysis.

#### Variable importance with explainable-AI (xAI)

The mathematical functions used in neural network models are unknown, but it would be useful to identify the covariates that contribute the most to explaining each species incidences. We therefore carried out an ‘explainable-AI’ (xAI) analysis, using the `R` package `flashlight 0.8.0` (Mayer, 2021). In short, for each environmental-covariate, we shuffled its values in the dataset and estimated the drop in explanatory performance on the training data. The most important covariate is the one that, when permuted, degrades explanatory performance the most (see Supplementary Information: Variable importance with explainable AI (xAI)).

#### Prediction and visualisation of species distributions

Finally, after applying the final model to the test dataset, we identified 76 species that had moderate to high predictive performance (AUC ≥ 70%). We used the fitted model and the environmental-covariates to predict the probability of each species’ incidence in each grid cell of the study area (‘filling in the blanks’ between the sampling points). The output of this one model is 76 individual and continuous species distribution maps, which we combined to carry out three landscape analyses. First, we counted the number of species predicted to be present (probability of presence ≥ 50%) in each grid square to produce a species richness map. Second, we carried out a dimension-reduction analysis, also known as ordination, using the T-SNE method (van der Maaten and Hinton, 2008; Krijthe, 2015) to summarise species compositional change across the landscape. Pixels that have similar species compositions receive similar T-SNE values, which can be visualised. Third, we calculated Baisero et al.’s (2022) site-irreplaceability index for every pixel. This index is the probability that loss of that pixel would prevent achieving the conservation target for at least one of the 76 species, where the conservation target is set to be 50% of the species’ total incidence.

Finally, we carried out *post-hoc* analyses by plotting site irreplaceability, composition (T-SNE), and species richness against elevation, old-growth structural index (Davis et al., 2015), and inside/outside HJA.

## RESULTS

### Model Inputs

#### DNA/Taxonomic data

The 121 samples from July 2018 were sequenced to a mean depth of 29.0 million read-pairs 150-bp (median 28.9 M, range 20.8-47.1 M). Of the 190 OTUs used in our joint species distribution model, 183 were assigned to Insecta, and 7 to Arachnida (Figure 1b). All OTUs could be assigned to order level, 178 to family level, 131 to genus level, and 66 to species level (Figures 1b, 4S).

### Statistical Analyses

#### Model performance and xAI

Across all species together, the final JSDM model achieves median and mean explanatory-performance values of *AUC* = 0.86 and 0.86, respectively, where the AUC (Area Under the Curve) metric equals 1 for a model with 100% correct predictions and 0 for 100% incorrect predictions. The model’s median and mean predictive AUC (i.e. on the test data) are 0.67 and 0.67 (Figure 2Sa). Predictive AUC is a measure of model generality, and the fact that explanatory AUCs are greater than predictive AUCs demonstrates how fitting a model to a particular dataset results in a degree of overfitting. Per species, mean AUC values range from 0 (fail completely) to 1 (predict perfectly), and this variation was not explained by species’ taxonomic family or prevalence (% presence in sampling points).

**Figure 2.**
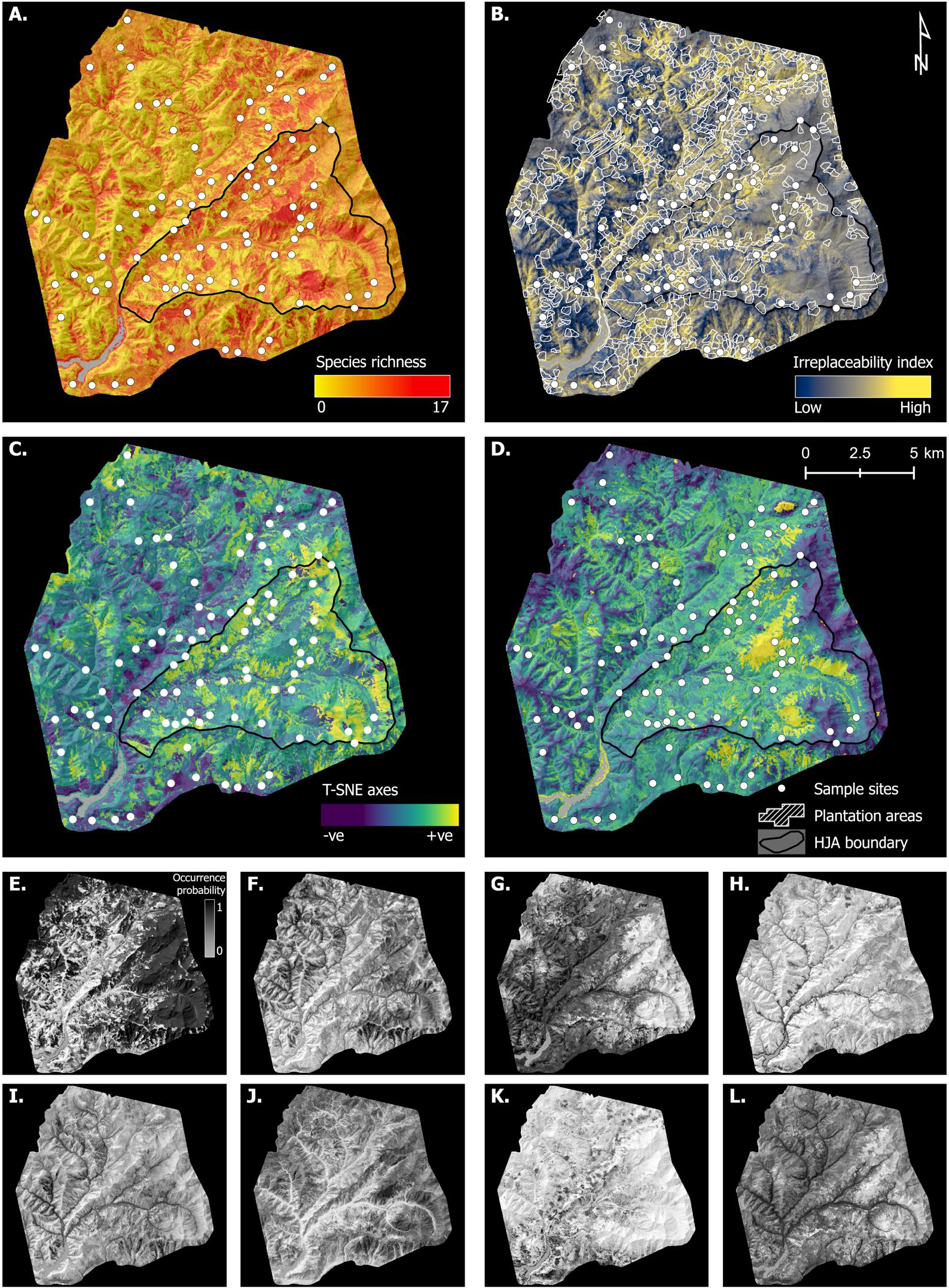
JSDM-interpolated spatial variation in species richness, irreplaceability, and composition, plus examples of individual species distributions. A. Species richness. B. Site beta irreplaceability, showing areas of forest plantation. C-D. T-SNE axes 1 and 2. White circles indicate sampling points, white polygons indicate plantation areas (i.e. a record of logging in the last 100 years), and the black-line-bordered triangular area delimits the H.J. Andrews Experimental Forest (HJA, Fig. 1). E-L. Selected individual species distributions (all species in Figure 9S), with BOLD ID, predictive AUC, and prevalence. E. Rhagionidae gen. sp. (`BOLD: ACX1094`, AUC: 0.91, Prev: 0.64). F. *Plagodis pulveraria* (`BOLD: AAA6013`, AUC: 0.81, Prev: 0.23). G. *Phaonia* sp.(`BOLD: ACI3443`, AUC: 0.80, Prev: 0.65). H. *Melanostoma mellinum* (`BOLD: AAB2866`, AUC: 0.90, Prev: 0.11). I. *Helina* sp. (`BOLD: ACE8833`, AUC: 0.73, Prev: 0.23). J. *Bombus sitkensis* (`BOLD: AAI4757`, AUC: 0.98, Prev: 0.23). K. *Blastobasis glandulella* (`Bold: AAG8588`, AUC: 0.86, Prev: 0.18). L. *Gamepenthes* sp. (`BOLD: ACI5218`, AUC: 0.77, Prev: 0.57).

Mean predictive AUC value does not increase with OTU abundance (as measured by incidence), and variability in predictive AUC values is only weakly higher in low-incidence OTUs (Figure 12S), especially for the OTUs with high mean predictive AUCs (i.e. those used to map species richness, composition, and site irreplaceability).

Out of 29 environmental covariates, 18 (Table 1S) were the most important for at least one species (Figure 2Sb). Elevation and Topographic Roughness Index (TRI) were the most important covariates for the most species. Eleven environmental covariates were the most important for at least one species in terms of interaction effects of the variables, with elevation and TRI again being the most important (Figure 8S).

#### Prediction and visualization of species distributions

Finally, we reduced the dataset to the 76 species with individual predictive AUCs ≥ 0.7 (mean = 0.834), and for each, we generated individual distribution maps across the study area, which differ in amount and distribution of the areas with high predicted habitat suitability (Figures 2 E-L, 9S). We then combined the maps to estimate the fine-scale spatial distributions of species richness, community composition, and site irreplaceability across the study area (Figure 2). Site irreplaceability, which is a core concept in systematic conservation planning, ranks each site by its importance to the “efficient achievement of conservation objectives” (Kukkala and Moilanen, 2013). In practice, high-irreplaceability sites tend to house many species with small ranges and/or species with large ranges that we wish to conserve a large fraction of, such as endangered species.

Greater species richness was predicted for areas without recent logging, especially within the northeast and southeast sectors of the H.J. Andrews Experimental Forest (HJA), on west-facing slopes, and in the south of the study area (Figure 2 A). A *post-hoc* analysis found a non-linear increase in species richness in the largest patches of old-growth forest, which are inside the HJA (Figure 3 A, B).

**Figure 3.**
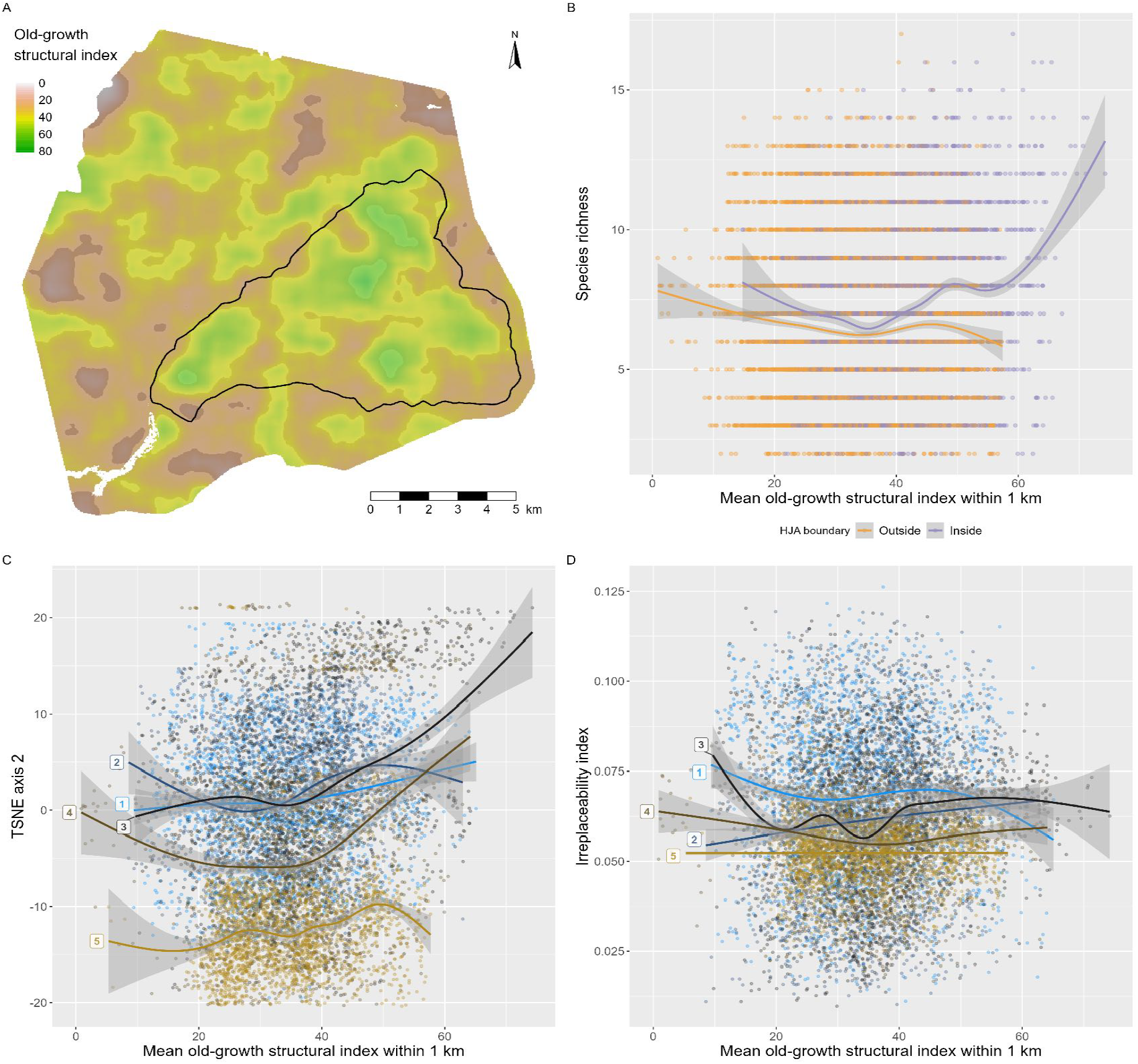
*Post-hoc* analysis of species richness, composition, and irreplaceability patterns in Figure 2, in relation to an old-growth structural index (OGSI) map, from Davis et al. (2015) A. Smoothed OGSI, showing principal patches of old-growth forest inside and outside the H.J. Andrews Experimental Forest (black-line-bordered triangular area). The HJA has the largest patches of old-growth forest. B. Species richness increases in the parts of the HJA with the highest OGSI values (compare with Figure 2 A). C. Species compositions in the largest old-growth patches, which are at elevation bands 3 and 4, are distinct from the rest of the landscape (compare with Figure 2 D). D. Irreplaceability shows no relationship with OGSI at any elevation (compare with Figure 2 B). Elevation bands (blue to brown colour gradient) 1: 380 − 620; 2: *>* 620 − 865; 3: *>* 865 − 1115; 4: *>* 1115 − 1365; 5: *>* 1365 − 1615 m above sea level. Splines fit using `mgcv` (Wood, 2017).

T-SNE ordination reveals spatial patterning in species composition (Figure 2 C, D). T-SNE-1 is clearly correlated with elevation (compare Figures 1a and 3 C), whereas T-SNE-2 (like species richness) appears to be correlated with the extent of surrounding old-growth forest, but only at middle elevations (Figure 3 C). Finally, site irreplaceability clearly follows stream courses, which are mostly at low elevations (Figure 2 B) and cover a small portion of the total landscape. As a result, *post-hoc* analysis also shows that irreplaceability decreases with elevation but finds no relationship between irreplaceability and surrounding old-growth forest (Figure 3 D).

## DISCUSSION

We combined *in-silico* barcode-mapping data derived from 121 arthropod bulk samples in 89 sampling points spread over a 225 km^2^ working and primary forest with 29 environmental covariates (Figure 5S) from Landsat, lidar, and other layers that covered information on forest structure, vegetation condition, topography, and anthropogenic impact. We used a joint species distribution model with a deep neural network to predict the fine-scale spatial distributions of 76 Insecta and Arachnida species with a high degree of estimated predictive performance (all individual predictive AUCs *>* 0.7, mean = 0.834) (Figure 2Sa). The model made good use of the 29 environmental covariates, with 18 of them being the most important for at least one species (Figure 2Sb), with elevation and Topographic Roughness Index (TRI) most important covariates for the most species. These two covariates were also the most frequently most important in terms of their interactions with other covariates (Figure 8S).

By interpolating to create continuous species distribution maps and combining them, we created *granular* maps of arthropod biodiversity metrics: species richness, community composition, and site irreplaceability (Figure 2). We observed *post-hoc* that species richness is higher and that species composition is distinct in the largest patches of old-growth forest (Figure 3 B, C), but not exclusively so. Irreplaceability, as we have defined it here using Baisero et al.’s (2022) formulation, which does not take connectivity or ecosystem functions into account, is highest along stream courses (Figure 3 D), which are dominated by species with high occurrence probabilities covering a small area (Figure 9S). Irreplaceability is not higher in old-growth forest, given that old-growth is not a rare habitat in our study area. We consider the patterns observed in (Figure 3) to be hypotheses for future testing, and thus we do not calculate statistical significance values.

A biodiversity map is more *understandable* than is an analysis of data points and can be compared directly with land-use maps. In principle, these datasets and products can also be *timely*, given that the creation of DNA-based datasets can be outsourced to commercial labs in some countries with turnaround times measured in weeks. Information *quality* can be assessed via prediction performance (Figure 2Sa), and even *trustworthiness* can be assessed via a combination of proof-of-work GPS surveyor tracking and independent re-sampling, given that sampling is standardized (Hartig et al., 2023).

In summary, we show how to generate information on arthropod spatial distributions with a high-enough resolution to make it useful and understandable for local management while also being efficient and standardised enough to scale up to thousands of square kilometres. However, as shown by the many species with low predictive AUCs (Figure 2Sa), future work will be needed to improve how error is accounted for when generating model outputs (Hartig et al., 2023; Diana et al., 2022), and we discuss methods for doing this in Supplementary Information: Caveats. We conclude by briefly reviewing potential applications of this approach.

### Potential applications of efficient, fine-scale, and large-scale species distribution mapping

This study demonstrates how the major steps of species distribution mapping are enjoying major efficiency gains (Besson et al., 2022; Bush et al., 2017; Speaker et al., 2022; Tosa et al., 2021). Large numbers of point samples can be characterized to species resolution via DNA sequencing and/or electronic sensors, large numbers of environmental covariates are available from near- and remote-sensing sources (Lock et al., 2022), and GPU-accelerated deep learning algorithms can be used to both accelerate and improve model fitting on these larger datasets (Pichler and Hartig, 2021, 2023). Although this study focused on arthropods, a wide range of animal, fungal, and plant taxa can be detected using DNA extracted from water, air, invertebrate, and soil samples (Abrego et al., 2018; Bohmann et al., 2014; Guimarães Sales et al., 2020; Ji et al., 2022; Leempoel et al., 2019; Lin et al., 2021; Massey et al., 2022; Rodgers et al., 2017; Thomsen and Sigsgaard, 2019; Tilker et al., 2020), with river networks being an especially promising way to scale up sampling over large areas (Guimarães Sales et al., 2020; Lyet et al., 2021).

As a result, it is possible to envisage implementing Pollock et al.’s (2020) vision of using ‘sideways’ species-based biodiversity monitoring to subdivide whole landscapes for ranking by conservation value (see also Cavender-Bares et al., 2022). One potential benefit would be to interpret remote-sensing imagery in terms of species compositions, thus improving the efficiency of habitat-based offset schemes, such as England’s Biodiversity Net Gain legislation, which has been criticized for undervaluing some habitat types, such as scrubland, that are known to support high insect diversity and abundance (Weston, 2021).

Recent studies have also shown that timely and/or fine-resolution biodiversity distribution data can potentially improve conservation decision-making, over that informed by historical distribution data. Ji et al. (2022) used 30, 000 leeches mass-collected by park rangers to map for the first time the distributions of 86 species of mammals, amphibians, birds and squamates across a 677 km^2^ nature reserve in China, finding that domestic species (cows, goats, and sheep) dominated at low elevations, whereas most wildlife species were limited to mid- and high-elevation portions of the reserve. Before this study, no comprehensive survey had taken place since 1985, impeding assessment of the reserve’s effectiveness, which is a general problem in the management of protected areas (Maxwell et al., 2020). Chiaverini et al. (2022) used camera-trap data to extrapolate the distributions of vertebrate species richness across Borneo and Sumatra and found that high species richness areas did not correlate well with IUCN range maps, which are based on historical distribution data (https://www.iucnredlist.org, accessed 18 April 2022). Finally, Hamilton et al. (2022) compiled decades of standardized biodiversity inventory data for 2216 species in the continental United States and interpolated to identify areas of unprotected biodiversity importance (using a measure similar to site irreplaceability, i.e. protection-weighted range-size rarity). Because the resulting maps were *granular* (990 m), Hamilton et al. (2022) were able to compare species distributions with land tenure data, including protected areas, and found large concentrations of unprotected species in areas not previously flagged in continental- and regional-scale analyses, in part due to the inclusion of taxa not normally included in such analyses (especially plants, freshwater invertebrates, and pollinators).

## Conclusion

A major difficulty for basic and applied community ecology is the collection of many standardised observations of many species. DNA-based methods provide capacity for collecting data on many species at once, but costs scale with sample number. In contrast, remote-sensing imagery provides continuous-space and near-continuous-time environmental data, but most species are invisible to electronic sensors. By combining the two, we show that it is possible to create a combined spatio(temporal) data product that can be interrogated in the same way as an exhaustive community inventory.

## ACKNOWLEDGMENTS

We thank field technicians BP Murley, SD Sparrow, and ME Yates. DWY and ML were supported by the Key Research Program of Frontier Sciences, CAS (QYZDY-SSW-SMC024), the Strategic Priority Research Program of Chinese Academy of Sciences, Grant No. XDA20050202, the State Key Laboratory of Genetic Resources and Evolution (GREKF19-01, GREKF20-01, GREKF21-01) at the Kunming Institute of Zoology, and the University of Chinese Academy of Sciences. DWY was also supported by the University of East Anglia and a Leverhulme Trust Research Fellowship (RF-2017-342), and benefited from the sCom Working Group at iDiv.de. MIT was supported by the National Science Foundation-funded H.J. Andrews Long-Term Ecological Research (LTER) program (DEB-1440409), Oregon State University, the ARCS Oregon Chapter, and the U.S. Department of Agriculture Forest Service. Field data collection was funded by Oregon State University, the Pacific Northwest Research Station, and the U.S. Department of Agriculture Forest Service. Lidar data processing was supported by the National Science Foundation-funded H. J. Andrews LTER Program (DEB-2025755, DEB-1440409) and the Pacific Northwest Research Station. The findings and conclusions in this publication are those of the authors and should not be construed to represent any official U.S. Department of Agriculture or U.S. Government determination or policy. The use of trade or firm names in this publication is for reader information and does not imply endorsement by the U.S. Government of any product or service.

## AUTHORS’ CONTRIBUTIONS

DWY and TL conceived the project. TL, MIT, DMB, DBL designed the sampling methodology; MIT and ML collected the data; YL, CD, ML, and DWY analyzed the data; PG and MP contributed unpublished software; YL, CD, and DY led the writing of the manuscript. All authors contributed critically to the drafts and gave final approval for publication.

## DATA AVAILABILITY

Raw sequence data are archived at NCBI Short Read Archive BioProject PRJNA869351. All scripts and data tables (from bioinformatic processing to statistical analysis to figure generation) are available at https://github.com/chnpenny/HJA_analyses_Kelpie_clean/releases/tag/v1.1.0 and archived at doi:10.5281/zenodo.8303158.

## COMPETING INTERESTS

DWY is a co-founder of NatureMetrics (www.naturemetrics.com), which provides commercial metabarcoding services. All other authors have no competing interests.

## Supplementary Information for the Article

### Dietz et al.’s five elements and the creation of a biodiversity offset market

A rare example of all five elements working together to achieve biodiversity conservation is the UK District Licensing offset market for the great crested newt (*Triturus cristatus*). Until recently, builders had been required to survey for the newt when their plans might affect ponds, and to respond to newt detections by paying for mitigation measures. Traditional surveys required at least four visits per pond during the short breeding season. After Biggs et al. [2015] showed that a single environmental-DNA (eDNA) water survey per pond, analysed with probe-based quantitative PCR (qPCR), could detect the newt with equal sensitivity (i.e. eDNA information is *high-quality* and *granular*), the UK government authorised newt eDNA surveys, and a private laboratory market grew to *provide the infrastructure* for *timely* and *trustworthy* information, via response times of a few days and an annual proficiency test. The switch to eDNA increased survey efficiency, but still left in place the UK’s reactive approach to newt conservation (‘mitigate after impact’). Mitigation measures, such as translocation, can delay building by over a year. In 2018, the UK government took further advantage of eDNA’s detection efficiency by implementing an *institutional redesign* with the District Licensing scheme, where hundreds of ponds across one or more local planning authorities are first systematically surveyed with eDNA [Natural England, 2019]. The data are used to fit a species distribution model, which is converted to an *understandable* map of discrete risk zones for the newt. Builders can now meet their legal obligations at any time by paying for a license, the cost of which depends on their site’s size, risk-zone level, and number of affected ponds, eliminating delay. The licence fees fund the proactive creation and long-term management of compensation habitat, including four new ponds per affected pond. Compensation habitat is directed toward Strategic Opportunity Areas, which reflect planning-authority building aspirations (*political bargaining*), and *enforcement* is through the same processes that apply to all planning permissions.

### Materials and Methods

#### Model Inputs

##### Field data collection

We collected 121 Malaise-trap samples of arthropods at 89 sampling sites in and around the H.J. Andrews Experimental Forest and Long-Term Ecological Research site (HJA), Oregon, USA in July 2018. Sites were stratified (as best as possible while yielding to logistical constraints) based on elevation and time since disturbance. Sites were also stratified between inside and outside the HJA to capture landscape-scale differences between a long-term ecological research site where no logging has occurred since 1989 and neighboring sites within a landscape context with continued active management. Each trap was left to collect for seven days, and samples were transferred to fresh 100% ethanol to store at room temperature until extraction. In 32 of the sites, two Malaise traps were set 40 m apart, and in the other 57, only one trap was set (Figure 1a). In August 2018, we repeated the sampling and processed all 242 samples together, but we have analyzed only the July samples for this study.

#### Wet-lab pipeline and bioinformatics

We follow the SPIKEPIPE protocol from Ji et al. [2020], where we map paired-end reads from Illumina shotgun-sequenced samples to a reference dataset of DNA barcode sequences. In shotgun sequencing, the total DNA of each sample is sequenced (the term shotgun refers to the random subset of the total DNA that gets sequenced), and the output ‘reads’ are ‘mapped’ (matched) to a reference set of barcodes. This approach relies on the enormous data output of Illumina sequencers, since only ∼ 1*/*4000 reads is from a DNA barcode, as opposed to the rest of the genome.

A major benefit of the SPIKEPIPE method is reduced workload since all that is needed is to extract DNA from each sample before sending to a sequencing center. The main disadvantage is that species present at low overall biomass are unlikely to be detected (although this is also a partial advantage in that any sample cross-contamination is also unlikely to be detected). However, low-biomass species are less likely to contribute meaningfully to species distribution modelling since the numbers of incidences for rare species are, by definition, low.

An important difference of this study from Ji et al. [2020] is that their study used a pre-existing reference set of DNA barcodes [Wirta et al., 2014], whereas we generate our reference set directly from the same shotgun-sequenced datasets, using the program Kelpie [Greenfield et al., 2019], which is an *in-silico* PCR program.

For this study, we only analyzed the July 2018 samples (*n* = 121), but the arthropod samples of both sessions were together extracted, sequenced, analyzed, and assigned to taxonomies.

#### DNA extraction and sequencing

Before extraction, we kept only the heads of insects with body sizes longer than 2 cm. DNA was non-destructively extracted by soaking the samples in 5X lysis buffer while shaking and incubating the samples at 56 ^*?*^C for 60 h [for more details, see Ji et al., 2020].

To the lysis buffers, we added a DNA spike-in standard of two beetle species in a 9 : 1 ratio. We shotgun-sequenced all 242 samples (PE 150, 350 bp insert size) to a mean depth of 29.0 million read pairs (range 21-47) on an Illumina NovaSeq 6000 at Novogene (Beijing, China). We used TrimGalore 0.4.5 (https://www.bioinformatics.babraham.ac.uk/projects/trim_galore, accessed 10 Sep 2021) to remove residual adapters (--paired --length 100 -trim-n).

#### Creating a barcode reference database using *Kelpie in-silico* PCR

In physical PCR, two specially designed DNA sequences known as PCR primers are used to amplify (make many copies of) a target sequence, which, here, is the portion of the mitochondrial cytochrome oxidase subunit I (COI) gene that is widely used as the taxonomically informative ‘DNA barcode’. If we had tried to use physical PCR to construct a reference library of DNA barcodes from the Malaise trap sample set, we would have needed to individually separate, sort, identify, extract, and PCR many hundreds of specimens.

Instead, we used a recently available shortcut known as ‘in-silico PCR’, using a software package called Kelpie [Greenfield et al., 2019]. Using the shotgun-sequence read files from the Malaise-trap samples, Kelpie carries out a computer search for reads that match the two ends of the target DNA barcode and then searches for overlapping reads, ultimately assembling DNA barcode sequences from the shotgun datasets. In our case, we use the BF3+BR2 primers from Elbrecht et al. [2019], which bookend a 418-bp fragment of the COI DNA barcode. After running Kelpie on all individual and groups of Malaise trap samples, `Kelpie` assembled 5560 unique DNA-barcode sequences, some more abundant than others.

We first used `FilterReads` to reduce the shotgun datasets to reads that resemble COI sequences (`FilterReads -qt 30 +f GenBank 24919 COI C99 20.mer 25pct input.fq`), using a reference kmer dataset `GenBank 24919 COI C99 20.mer` (accessed 3 Aug 2021). This step is optional but greatly increases efficiency. We then used `Kelpie 2.0.11` [Greenfield et al., 2019] to carry out *in-silico* PCR on the filtered datasets (`Kelpie -f CCHGAYATRGCHTTYCCHCG -r TCDGGRTGNCCRAARAAYCA -primers -filtered -min 400 -max 500`). Binaries for both are at https://github.com/PaulGreenfieldOz/WorkingDogs/tree/master/Kelpie_v2 (accessed 20 Nov 2023). Kelpie mimics PCR on shotgun datasets by finding reads that include the forward primer sequence and step-by-step overlapping reads until a read matching the reverse primer is found. The advantages are that it is trivial to switch primers, lab workload is reduced, there can be no PCR error or PCR contamination, and the primer regions are returned.

The main disadvantage of *Kelpie* is that low-abundance species in a sample are usually not detected since every species requires enough reads in the dataset to complete the assembly from the forward to the reverse primer. That said, low-biomass OTUs are unlikely to contribute much to modelling, as they are also likely to exhibit low prevalence (few detection events) in the dataset. Nonetheless, we still tried to retrieve as many OTUs as possible by running *Kelpie* individually on each of the 242 samples and also running on concatenated fastq files made up of sample clusters (each site and its five nearest neighbors). The logic for the two steps is that even rare species might be abundant somewhere. In our experience, it is not helpful to concatenate large numbers of sequence files because rare amplicons look like error variants when there also exists in the dataset a similar but abundant amplicon sequence. *Kelpie* removes such rare amplicons as part of its error correction procedure. We combined the *Kelpie* outputs, gave the sequences unique names, and dereplicated, resulting in 5560 unique sequences.

The variation represented by these 5560 unique sequences derives from multiple causes: true genetic differences among species, true genetic diversity within species, errors generated by the Illumina sequencer, and rare pseudogene sequences from mitochondrial DNA that got copied into the nuclear genome at various points in each species’ past and been released from purifying selection. The latter are known as NUMTs (nuclear mitochondrial DNA).

We assigned taxonomies to all 5560 unique sequences on https://www.gbif.org/tools/sequence-id (accessed 3 Aug 2021), which provides three sequence-match classes (`‘exact’, ‘close’`, and `‘no’` match). For the exact match class, we retained the assignment to species, for the close match class, we retained the assigned genus and used NA for the species epithet, and for the weak match class, we retained the assigned order and used NA for lower ranks. We deleted all sequences that received a `‘no match’` or were not assigned to Insecta or Arachnida, after which, we used `vsearch 2.15.0` to cluster the sequences into 1538 97%-similarity OTUs.

Although PCR error has been avoided, Kelpie amplicons unavoidably still include Illumina sequencer error, including homopolymers (incorrect nucleotide repeats), which induce frameshift mutations. However, because the amplicon is of a protein-coding gene, we aligned the OTU representative sequences by their inferred amino-sequences (‘translation alignment’), using the invertebrate mitochondrial code in `RevMet 2.0` [Wernersson, 2003], after which we curated the sequences by eye, fixing obvious homopolymer errors and removing sequences with uncorrectable stop codons and those that failed to align well to the others, the latter two likely being ‘Numts’ (pseudogenes from nuclear insertions of mitochondrial sequences). This left us with 1520 OTUs.

In the final step, we read in the taxonomies of these OTUs and visually checked pairs of OTUs that had received very similar taxonomies (ID’d to the same BOLDID) for which one OTU contained many reads and the other contained few. These are likely oversplit OTUs, and we removed the smaller of the OTUs. In rare cases, there are multiple OTUs that match to the same BOLDID, but one or more of them are only BLAST weak matches to that BOLDID and contain many reads, suggesting that these OTUs are true species for which reference sequences do not exist. Our bias throughout is to remove OTUs that could be artefactual splits of true OTUs, because these small OTUs will interfere with read mapping and do not add true diversity to the dataset. We were left with 1225 OTUs as the reference barcode set, and to this fasta file, we added the two spike-in COI sequences.

#### Read mapping with minimap2, samtools, and bedtools

We then used the newly constructed reference barcode dataset to detect species in each sample’s shotgun reads. This is done by applying a commonly used tool from genomics known as a sequence alignment program, which maps individual Illumina reads against one or more reference sequences (usually a genome, but here the reference barcodes). Reference barcodes to which multiple Illumina reads are aligned are taken to be present in that sample, as long as the read mappings are (1) high quality (close match, low estimated error rate, map in the correct orientation) and (2) cover more than 50% of the barcode length, under the logic that if a species is truly in a sample, reads from the whole COI gene will be in the sample and will thus ‘map’ along the length of that species’ barcode. These acceptance criteria were determined with experimental mock samples of known composition [Ji et al., 2020]. The output of mapping all samples individually to the reference barcodes is a sample x species table. After removing a few samples that were missing sample-identifying metadata or had no mapped reads to the spike-ins, we were left with 237 samples of the original 242, of which 121 were from sampling session 1 (July 2018).

We used `minimap2 2.17-r941` (Li 2018) in short-read mode (`minimap2 -ax sr`) to map the read pairs from each sample to the 1225 reference barcodes and the 2 spike-in sequences. We used `samtools 1.5` [Li, 2018] to sort, convert to bam format, exclude reads that were unmapped or mapped as secondary alignments and supplementary alignments, and include only ‘proper-pair’ read mappings (mapped in the correct orientation and at approximately the correct distance apart) at ≥ 48 ‘mapping quality’ (MAPQ) (`samtools view sort -b -F 2308 -f 0x2 -q 48`).

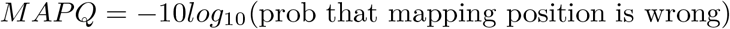

We accepted *MAPQ* ≥ 48 after inspection of the highly bimodal distribution of quality values, with most reads giving *MAPQ* = 60 (probability of error = 0.000001) or 0 (i.e. maps well to multiple locations). *MAPQ* = 48 corresponds to an error probability ∼ 0.000016. Informally, we have found that limiting quality to only the highest value, 60, has little effect on the results, whereas including low-quality mappings (`-q 1`) leads to more false-positive hits (data not shown). Read mapping data were output to `samtools idxstats` files.

The output for each sample is the number of mapped reads per OTU and spike-in that have passed the above filters. However, it is still possible for a barcode to receive false-positive mappings. Thus, we applied a second round of filtering. We expect that if a species is truly in a sample, reads from that sample will map *along the length* of that species’ barcode, resulting in a high percentage coverage. In contrast, if reads map to just one location on a barcode, even at high MAPQ, the percentage coverage will be low, and we consider those mappings to be false-positive detections caused by that mapped portion of the barcode being very similar to a species that is in the sample but not in the reference database. We used `bedtools 2.29.2` [Quinlan and Hall, 2010] to calculate the number of overlapping reads at each position along the reference sequence (`genomecov -d`). The percent coverage is the fraction of positions in a barcode covered by one or more mapped reads. We kept only those species detections with percent coverage ≥ 50%, following recommendations from an experiment in Ji et al. [2020].

#### Sample X Species table creation

We imported the sample metadata and the samtools and bedtools outputs into R 4.0.4 [R Core Team, 2022] for downstream processing into a sample x OTU table. After removing a few sites that had missing sample-identifying metadata or had no mapped reads to the spike-ins, we were left with 237 samples out of the original 242. These samples represented two sampling sessions, of which 121 were in sampling Session 1 (July 2018) and 116 in Session 2 (August 2018). The 121 samples from Session 1 were distributed over 89 sites, of which 57 sites had 1 Malaise trap-sample and 32 sites had 2 samples. For this study, we used only the Session 1 samples. The two sessions only partially overlapped in species composition, meaning that it was not possible to test a Session 1 model on Session 2.

#### Environmental covariates

We used environmental covariates related to forest structure, vegetation reflectance and phenology, topography, anthropogenic features, and location to model arthropod incidence. We extracted the forest structure variables from lidar data collected from 2008 to 2016, consisting of 95^th^ percentile canopy height, canopy cover above 2 and 4 m (calculated as the proportion of returns for a 30 m pixel above that height) and proportional area with canopy cover (calculated as the proportion of area with vegetation greater than 4 m) (Table 1S). These types of measures of canopy height and cover are correlated with field observations of forest structure in Pacific Northwest coniferous forests, such as mean diameter, canopy cover, and tree density [Kane et al., 2010]. We calculated vegetation indices from Landsat 8 images over the year, 2018, including Normalized Difference Vegetation Index (NDVI), Normalized Difference Moisture Index (NDMI), and Normalized Burn Ratio (NBR). From these, we calculated annual metrics of standard deviation, median, 5% and 95% percentiles over the year 2018, as well as using raw bands from a single cloudless image from 26/07/2018 (within 7 days of data collection). Both the proportion of canopy cover and annual Landsat metrics were calculated within the radii of 100, 250 and 500 m, given that vegetation structure at different spatial scales is known to drive arthropod biodiversity [Müller et al., 2014]. We created topographic predictors based on 1 m resolution bare-earth models from lidar ground returns, including elevation, slope, Eastness and Northness split from aspect, Topographic Position Index (TPI), Topographic Roughness Index (TRI) [Wilson et al., 2007], Topographic Wetness Index (TWI) [Metcalfe et al., 2018], and distance to streams, based on a vector stream network (http://oregonexplorer.info, accessed 24 Oct 2019). We used spatial data on anthropogenic activities to create predictors based on distance to nearest road, proportion of area logged within the last 100 and 40 years within radii of 250, 500 and 1000 m, and a categorical variable of inside or outside the boundary of the H.J. Andrews Experimental Forest. We used the raster and sf packages for R for all spatial analysis [Hijmans, 2022, Pebesma, 2018]. We mapped all 58 candidate environmental covariates (Table 1S) at 30 m resolution — either matching native resolution (e.g. Landsat), or aggregated from finer resolution data (e.g. lidar data), and projected them to the UTM 10N grid.

### Statistical Analyses

#### Species inputs

For modelling, we converted the sequence-read-number OTU table to presence-absence (1*/*0), and we only included OTUs present at ≥ 6 sampling sites across the 121 samples. Our species dataset thus consisted of 190 OTUs in two classes, Insecta and Arachnida (Figure 1b).

#### Environmental covariates

To avoid collinearity, which would pose problems for the application of explainable AI [xAI, see below; Hooker et al., 2021], we iteratively calculated the Variance Inflation Factor [VIF; Zuur et al., 2007] on the 58 scaled candidate covariates, eliminating the highest scoring variable each time until all VIF values were *<* 8. The exception is that we forced the covariates elevation and inside/outside H.J. Andrews Forest to remain within the set of predictors irrespective of their VIF value, for a total of 29 predictors.

#### Joint Species Distribution Model

The general idea behind species distribution modelling is to “predict a species’ distribution”, using the species’ observed incidences (presences and absences) and the combination of environmental-covariate values (i.e. the 29 covariates) in those points, to estimate the probability of species’ incidences (i.e. to ‘fit the model’). After model fitting, species in the rest of the sampling area, where environmental conditions are known but species’ incidences are not, can be predicted, and the fitted model uses the environmental-covariate values to calculate the species’ probability of presence. In this way, each species’ distribution is predicted across continuous space, with varying degrees of accuracy.

We used the R package sjSDM 1.0.5 [Pichler and Hartig, 2021], which is a JSDM that implements an integral approximation of multivariate probit models. sjSDM also includes a DNN (deep neural network) option to fit environmental covariates, which suits our dataset of many species with few data points and many covariates. We modeled the presence-absence data with a binomial distribution (probit link) in the sjSDM framework. The species occurrence probabilities are described as a function of a three-layer DNN on the environmental covariates in addition to spatial coordinates to account for spatial auto-correlation and a species covariance matrix:

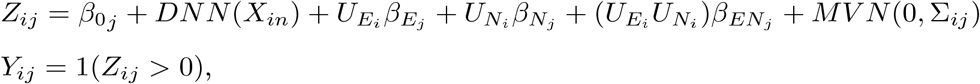

in which *Z*_*ij*_ is the occurrence probability of species *j* at sampling site *i*; *Y*_*ij*_ is the observed presence of species *j* at site *i*; *X*_*in*_ is the value of environmental covariate n in sampling site *i*. The second part of the model describes the trend-surface model, which is one way to account for spatial auto-correlation [Dormann et al., 2007]: 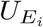 and 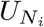 are the two Universal Transverse Mercator variables (coordinates) which are modeled for each species *j* at sampling site *i* as linear terms with coefficients 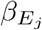 and 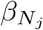, and as interaction with coefficients 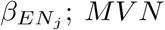 is the multivariate normal error representing the species correlation matrix.

#### Tuning and Testing

The statistical challenge is to avoid overfitting, which is when the fitted model does a good job of predicting the species’ incidences in the sampling points that were used to fit the model in the first place but does a bad job of predicting the species over the rest of the landscape. Overfitting is most likely to occur with species that have few presences, with large numbers of environmental covariates, and when the model uses flexible mathematical functions to describe the relationships between environmental-covariates and species incidences. Unfortunately, all three of these conditions apply when trying to model arthropod fine-scale distributions. Many species are rare, there are many candidate remote-sensing covariates, and we expect that any relationships between remote-sensing-derived covariates and arthropod incidences will be indirect and thus complex, necessitating the use of flexible mathematical functions.

To minimise the risk of overfitting, we applied a combination of regularisation and cross validation. Regularisation is a statistical method that reduces small (or uncertain or collinear) covariate effects to zero. In this way, the initially high complexity of a DNN algorithm can end in a DNN model with a low effective complexity with good generality, even for small data.

In Figure 1SB_2_, we list nine model ‘hyperparameters’, which consist of the weighting between lasso and ridge regularisation (*α*_*e,s,b*_) and their strengths (*λ*_*e,s,b*_) for each of the environmental, spatial, and species covariance components, plus the dropout rate, the hidden structure for the DNN, and the learning rate of the model (Figure 1SC). These hyperparameters govern the neural network’s structure and how it is fit to the data, and the challenge in fitting is to select optimal regularisation values (the alphas and lambdas, and the dropout rate) for accurate prediction, which we do via 5-fold, nested cross-validation, in a procedure known as model tuning.

First, we randomly split the 121 data points from July 2018 into 75% *training* data (*n* = 91) and 25% *test* data (*n* = 30) (the latter also known as hold-out data, or outer split), and we ensured that when two Malaise traps had been placed at the same site, they were assigned to the same split (Figure 1SA).

We then worked with only the *training* dataset for model tuning (i.e. inner split). We split the training dataset into five ‘folds’ (=sections), also ensuring that data from pairs of traps placed at the same site were assigned to the same fold. We chose one combination of hyperparameter values, fit the model with those hyperparameter values to 4 of the 5 folds (as a single dataset), and measured how well this fitted model predicted presences and absences in the sites from the fifth fold (the *validation* dataset), which the model had *not* been fit to. This is the model’s predictive performance on that fold with that hyperparameter combination. Because we chose **5**-fold CV, we repeated this procedure five times, each time predicting a different fold of the five (Figure 1SB_1_). We calculated the model’s mean predictive performance over the five validation datasets and the model’s mean explanatory performance on the five training datasets. We repeated this five-fold CV procedure for 1000 hyperparameter combinations sampled from the total set of possible hyperparameter combinations (*n* = 7200), recording all 1000 mean performances in Figure (1SC) (black pts: mean predictive performances. blue pts: mean explanatory performances). We used six metrics to evaluate predictive and explanatory performance: AUC (area under the receiver operating characteristic curve), positive likelihood ratio, Pearson’s correlation coefficient, log-likelihood, True Skill Statistic (TSS), and Nagelkerke’s *R*^2^ [Lawson et al., 2014, Wilkinson et al., 2021, see Supplementary Information].

From the set of 1000 models, we chose the model with the hyperparameter combination that produced the highest predictive performance (designated as the tuned model) and fit it to the full training dataset (i.e. no folds). This is the *Final fit* model (Figure 1S*B*_1_), which we used to calculate explanatory performances per species. Finally, we also used *Final fit* model to predict presences/absences in the 25% *test* dataset that the model had never seen and calculated a predictive performance per species: AUC_pred_. The final models chosen by the other performance metrics behaved similarly (Figure 3S).

For species mapping, we filtered to those species that showed moderate to good predictive performance (*AUC*_*pred*_ *>* 0.70, *mean* = 0.83). **The key point is that because AUC**_**pred**_ **is calculated from the test dataset, *which the model never saw during tuning and final fitting***, **we can use each species’ AUC**_**pred**_ **as a measure of model generality for that species, and high AUC**_**pred**_ **species are therefore the species for which overfitting is a low risk**.

The purpose of regularisation is to create simpler, but not too simple, models, and this has the effect of creating models that are more likely to be general. When using regularisation, one is freed to use large numbers of covariates and terms in the model because regularisation typically sets most of their coefficients to 0. **The cost of regularisation is that one needs a large number of samples for model tuning (selecting the optimal regularisation regime via cross-validation) and to provide an untouched dataset for measuring model predictive performance**.

#### Variability in Predictive AUC by OTU Incidence

Finally, using a single holdout dataset for final testing does not allow estimates of the *variability* of our model (with respect to predictive AUC). We therefore ran an alternative model evaluation, using 5-fold cross validation over the whole dataset, which allows such an estimate. In this alternative evaluation, we followed the above methods to perform 5-fold cross validation, but now we used the entire dataset (121 points, 225 OTUs), with the same 75% - 25% splits for the training and validation folds, the same number of runs (1000 different combinations of hyperparameters), and the same prevalence threshold (minimum presence at 6 or more sites). From these runs, we chose the model with the highest predictive performance, as measured by AUC only in this case, and then used these hyperparameters to fit a model on the full dataset, producing the alternative *Final fit* model. Using this *Final fit* model, we ran a further 5-fold cross validation (using a 75%-25% split) and saved the results from each of the five validation predictions for all OTUs. The accuracy metrics were then averaged for each OTU and displayed graphically (Figure 12S). We ran a polynomial regression to test whether the standard deviation of predictive AUCs is greater for lower-incidence OTUs (Figure 12S). Finally, we used the OTUs with *AUC*_*pred*_ ≥ 0.70 (*n* = 112, *AUC*_*mean*_ = 0.80) to create maps of species richness, ordination axes, and irreplaceability (Figure 13S), in the same way as the main analysis.

We use this alternative analysis for the *sole purpose* of estimating the *variability* of predictive AUCs because fitting a model to the whole dataset increases the risk of overfitting and could potentially overestimate the predictive performance of the model, which is what we avoided by using a pure holdout in the main analysis. Ultimately, with a much larger dataset, running a nested CV with an inner k-CV (for training) and an inner k-CV (for testing) would be the gold standard to produce reliable estimates of the predictive AUCs and their variabilities together. However, given that typical ecological community datasets have many rare and many abundant species, splitting the data twice sequentially would likely frequently produce training and test splits with either no occurrences (for rare species) or only occurrences (for abundant species), making it technically impossible to fit reliable models or to validate them fairly. This suggests that the question of how to effectively evaluate and tune ecological community models should continue to be a priority for future research.

**Figure 1S.**
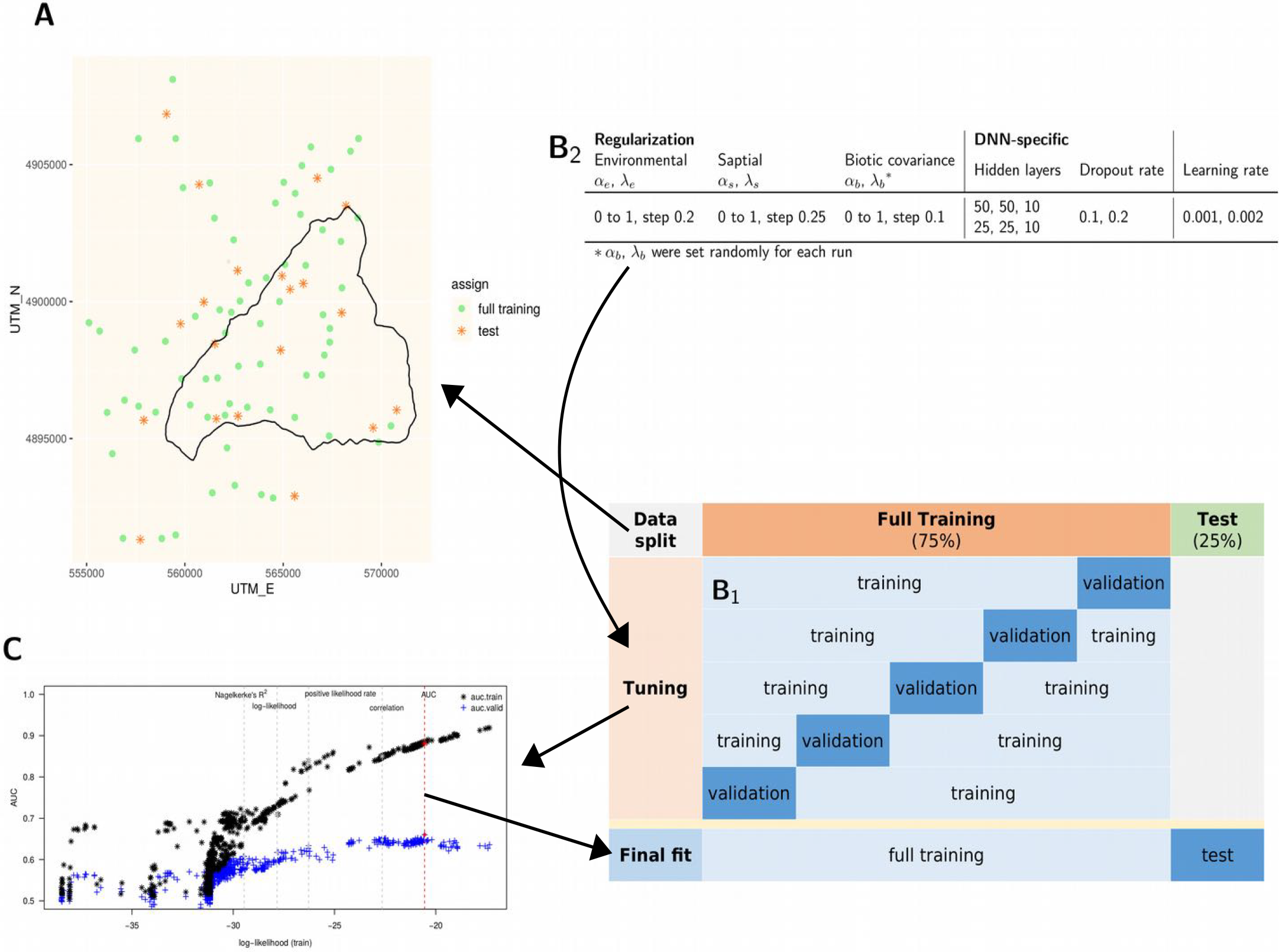
Model tuning and training strategy. We obtained our final model by data splitting, tuning, and final fitting. *A*. We randomly split the 121 Malaise traps into test (*n* = 30) and training subsets (91). *B*_1_. We then randomly split the training set into five parts for tuning via a 5-fold cross-validation. For all sets of splits, when a sampling site contained two Malaise traps, both traps were assigned to the same split. During each round of tuning (same hyperparameters combination), five models are run with one fold as the validation data and four folds as training. *B*_2_. We randomly sampled 1000 rows from a tuning grid of all combinations of hyperparameters (*n* = 7200), and the performance of each tuning model was tested against the validation data. *λ* sets the overall strength of regularization, and *α* sets the relative weighting of ridge vs. lasso penalties. *C*. After finding the best combination of hyperparameters for the AUC (area under the ROC curve) performance metric, we fit the model to the full training data and tested the fitted model’s predictive power against the test data. The black asterisks are the average AUC values for the training sets, and the blue crosses are the average for the validation sets.

**Figure 2S.**
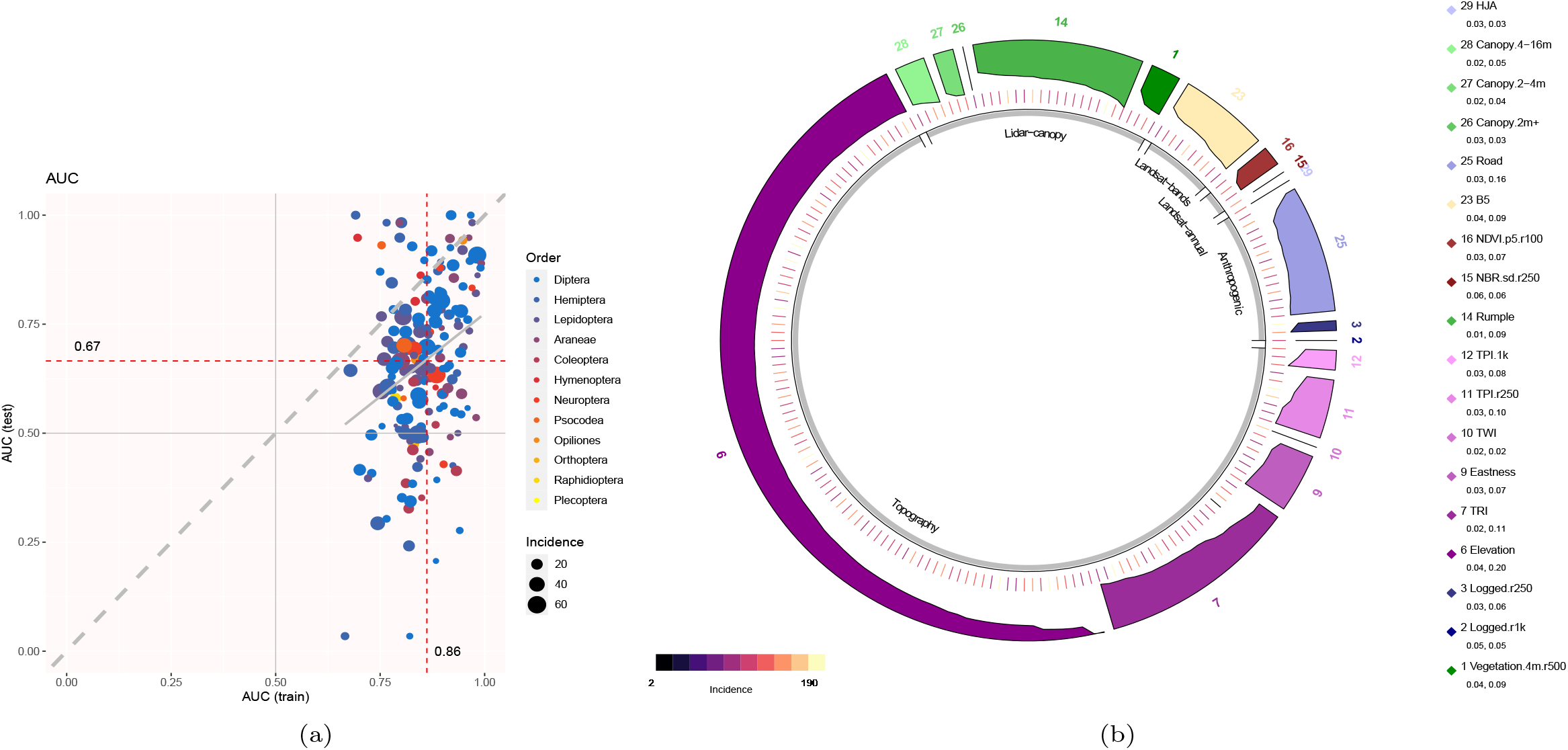
Model performance and environmental-covariate importance. (a). Explanatory AUC (range 0.67-1, mean 0.86, median 0.86) and predictive AUC (range 0.03-1, mean 0.67, median 0.67) of the final model. Each point is one OTU. Color indicates taxonomic class (order), and point size indicates incidence (number of Malaise traps in which the OTU was detected). Predictive AUC value is not explained by incidence (linear model, p = 0.93, R^2^ = 4.5*e* − 05). The dashed gray line is the 1:1 line, and the solid gray line is a fitted linear regression. (b). Most important explanatory environmental covariate for each OTU, as determined by xAI (see Variable importance with explainable AI). Tick marks indicate each OTU’s incidence, color bands indicate individual covariates, and gray bands indicate logical covariate groupings (Table 1S). Elevation (variable 6) and Topographic Roughness Index (variable 7) are the most important individual environmental covariates for the most OTUs, and the six variables in the topography group are the most important as a group. The heights of the colour bars are scaled to the permutation importance for that OTU.

#### Variable importance with explainable AI (xAI)

To gain insight into the importances of the environmental covariates in our DNN, we analyzed variable importance using permutation and Friedman’s H statistics, as implemented in the R package `flashlight 0.8.0` [Maksymiuk et al., 2020, Mayer, 2021].

Variable importance is based on global permutations of variables in the dataset [Fisher et al., 2019]. The calculation consists of several steps: First, a variable *x*_*i*_ from the dataset X is permuted (the values are randomised globally (over all sites)) and replaces the original *x*_*i*_ in *X*, so that we get a new dataset *X*_*permuted*_ with *x*_*i,permuted*_ (all other variables are not permuted). By permuting the variable, the effect (or association) between *x*_*i*_ and the response variable (e.g. species occurrence) is removed. Second, we generate new predictions with our model and dataset *X*_*permuted*_. Third, we calculate the predictive performance for our new predictions (here, AUC, see below). Fourth, we compare the new predictive performance for *X*_*permuted*_, which contains the permuted variable *x*_*i,permuted*_, with the predictive performance of the non-permuted dataset X. The difference between these two performances corresponds to the permutation importance of the variable *x*_*i*_. If *x*_*i*_ has a strong effect on the response variable, the permutation importance of variable *x*_*i*_ will be large because the model cannot predict the response well anymore. All these steps are repeated for all variables in the dataset. The advantage of this variable-importance protocol is that it does not require re-fitting the model n times for the n variables in the data set. We omitted the spatial component when calculating variable importance.

Friedman’s H-statistic is used to infer the importance of variable-variable interactions [Friedman and Popescu, 2008]. The statistic is based on partial dependencies (*PD*). Partial Dependencies describe the marginal effects of a variable on the response variable. Friedman’s H statistic additively decomposes the predict function 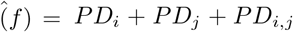, assuming that it consists of main effects (*PD*_*i*_ and *PD*_*j*_) and an interaction *PD*_*i,j*_ of two variables. Friedman’s H statistic estimates the importance of an interaction by comparing the interaction PD with the individual PDs: *PD*_*i,j*_ − *PD*_*i*_ + *PD*_*j*_. Without the subtraction, the interaction (*PD*_*i,j*_) would accumulate the individual effects (we only want the “shared” part). Finally, the variance of *PD*_*i,j*_ − *PD*_*i*_ + *PD*_*j*_ divided by the variance of *PD*_*i,j*_ corresponds to interaction importance between *x*_*i*_ and *x*_*j*_.

We calculated these xAI metrics based on the explanatory performance of the JSDM model, and the AUC performance matrix was used. The variable importance was calculated by permuting all data points of the environmental covariates over six repetitions to ensure a stable result. Afterwards, we chose the ten most important covariates based on the resulting variable importance for each species to conduct the unnormalized H-statistics. The unnormalized H-statistics were chosen to ensure a fair comparison between variables. The H-statistic was calculated using all the data points as well.

#### Prediction and visualisation of species distributions

Using the final model, we show three examples of how to visualize species predictions. Firstly, we used the final model to predict the distributions of those species with predictive AUC *>* 0.7. To avoid extrapolation [Norberg et al., 2019], we restricted predictions to a 1 km buffered, convex hull around all sample sites, edited manually to avoid suburban areas in the southern extreme of the study area. Further, all predictors within this area were restricted, or ‘clamped’, to lie within the range of predictor values across all sample points, that is, predictors above or below this range were given the maximum or minimum value from across the sample points, respectively [Anderson and Raza, 2010]. Given the stochasticity inherent in JSDM predictions based on sjSDM [Pichler and Hartig, 2021], each species’ prediction used the average of five separate prediction runs. We created binary species distributions maps by applying a 0.5 threshold on the occurrence probability values, and summed these to create a species richness map. We acknowledge that a common threshold for all species is not ideal, but no further analysis is performed with the binary maps.

Secondly, to map community similarity across the study area, we ordinated species predictions on two dimensions using T-SNE (t-Distributed Stochastic Neighbor Embedding) and mapped the two resulting ordination axes. T-SNE is a dimension-reduction technique where high-dimensional distances between data points are converted into conditional probabilities that represent similarities [van der Maaten and Hinton, 2008]. The R implementation [Krijthe, 2015] uses the Barnes-Hut approximation to increase performance with large data sets. The perplexity parameter, which controls the number of points available within the neighborhood, was set at 50.

Finally, after applying the final model to the test dataset, we identified 76 species that had moderate to high predictive performance. We used the fitted model and the environmental-covariates to predict the probability of each species’ incidence in each grid cells in the study area (‘filling in the blanks’ between the sampling points). The output is 76 individual and continuous species distribution maps, which we combined to carry out three landscape analyses. First, we counted the number of species predicted to be present (probability of presence ≥ 50%) in each grid square to produce a species richness map. Second, we carried out a dimension-reduction analysis, also known as ordination, using the T-SNE method [van der Maaten and Hinton, 2008, Krijthe, 2015] to summarise species compositional change across the landscape. Pixels that have similar species compositions receive similar T-SNE values, which can be visualised. Third, we calculated Baisero et al. [2022] site-irreplaceability index for every pixel. This index is the probability that loss of that pixel would prevent achieving the conservation target for at least one of the 76 species, where the conservation target is set to be 50% of the species’ total incidence.

Thirdly, we calculated the Baisero et al. [2022] site-irreplaceability index (*β*) per pixel across the study area as the combined probability that a site is irreplaceable for at least one OTU. The beta index combines species-level irreplaceability indices, alpha, at each site, measured as proximity-based metrics of how close a site is to being required to achieve a conservation target for a particular species. We used a value of 50% of each species’ total incidence across the study area as our conservation target.

Finally, we carried out post-hoc analyses by plotting site irreplaceability, composition (T-SNE), and species richness against elevation, old-growth structural index [Davis et al., 2015], and inside/outside HJA. We consider these analyses to be post-hoc because we are applying them to the predicted species distributions, which we viewed before analysis. Thus, we consider these analyses to be hypothesis-generating exercises for future studies.

### Caveats

#### Irreplaceability

We used Baisero et al.’s (2022) method to calculate site irreplaceability. Two advantages are that it is fast to calculate and is stable to changes in the grid system and in the addition or subtraction of species from the dataset, unlike the alternative method of using selection frequency from the outcome of a systematic conservation planning (SCP) algorithm, which must assume that the sites selected by any given SCP run are optimal. As Langford et al. [2011] point out, SCP algorithms are not widely tested for robustness to input error.

In contrast, Baisero et al.’s (2022) site-irreplaceability value is directly calculated: defined as one minus the probability that a site is replaceable for all species in that site. A value of 0 means that a site’s loss would still allow the conservation target of every species in that site to be met using other sites in the landscape, where a target is the proportion of a species’ range that is designated for protection. Thus, sites with higher irreplaceability values are characterised by higher numbers of species with high targets and/or small ranges. The latter reason is why lower elevations, the riverine basin (including the southern edge, which borders a river), and plantations are given high irreplaceability values (Figure 2 B), since these habitat types (and their associated species) cover a smaller proportion of the total landscape, and thus any species limited to them needs those sites protected for their conservation targets to be met (Figure 2 A). It is important to keep in mind that any measure of site irreplaceability can only compare the sites *within* the analysed landscape, meaning that a small pine plantation in a tropical rainforest would be scored high on irreplaceability if it contained pine-specialist arthropods. For such situations, known widespread and common species could be given low conservation targets, and artefactually rare habitats (the plantation in a rainforest) could be masked from analysis. For instance, we repeated the site-irreplaceability analysis after masking plantations, since recently logged forest characterises most of the Oregon forest landscape outside the H.J. Andrews Experimental Forest. Without plantations, areas near streams increased in irreplaceability value (Figure 10S).

Finally, given the rapidity with which Baisero et al.’s site-irreplaceability values can be calculated, one possible approach to account for error in predicted species distributions (Figure 9S) would be to resample the site-irreplaceability calculations in some way and to plot mean or median site irreplaceability values. The idea would be to produce a map that upweights the contributions of species with higher values of predictive performance and with higher occupancy probabilities. However, this proposed approach would require testing to see whether it in fact produces a more reliable map.

#### False-negative and false-positive errors

Despite detecting 1225 OTUs across the whole dataset, ultimately, only 190 OTUs had ≥ 6 detections. An independent analysis of this dataset has estimated that even the 50 most prevalent species have only a ∼ 50% probability of being detected when they are truly at the sampling points [Diana et al., 2022]. Consequently, we infer that many species absences are false negatives, which biases species prevalences and environmental-covariate effect sizes downwards. To increase the number of species that can be modelled, we make four recommendations:

1. Per sample, increase DNA-sequencing depth and/or increase the concentration of DNA barcode sequences using hybridisation or physical PCR [e.g. Liu et al., 2016, Yang et al., 2021].
2. Change the trapping method. Malaise traps seem especially prone to false-negative error [Steinke et al., 2021]. An alternative is pitfall traps, for which it is cheap to increase trapping effectiveness [by adding cups and guidance barriers, Boetzl et al., 2018].
3. Increase the number of sampling points. This would allow the training and test dataset sizes to be increased, allow more folds in the cross-validation step, and reduce the metrics of predictive performance, since AUCpred variance decreases with incidence.
4. Take multiple replicates per sampling point. Roughly, the per-bulk-arthropod-sample cost of the mitogenome mapping protocol is ∼ US$250, and commercial bulk-sample metabarcoding prices (i.e. physical PCR) range from US$100 to $350 per sample. Two traps per 89 sites would cost $17, 800 to $62, 300 total, or $79 to $277 per km^2^. Using multiple traps per site directly reduces the rate of false negatives, allows one to increase the minimum incidence threshold for inclusion in the model, and provides the option of combining occupancy correction and JSDMs [Doser et al., 2022, Tobler et al., 2019, Diana et al., 2022] to account for false-negative error.

#### Environmental covariates

We used both LANDSAT and multiple lidar datasets collected from 2008-2016 to generate predictors for species data collected in 2018, following successful use of Earth Observation data for biodiversity mapping in other studies [Bae et al., 2019, Galbraith et al., 2015, Lin et al., 2021, Müller et al., 2009, Müller and Brandl, 2009]. The temporal mismatch between lidar and field data might introduce some errors [Gatziolis and Andersen, 2008] if major vegetation changes had occurred between acquisitions (e.g. tree mortality), but in most cases, we expect forests to change slowly [Zald et al., 2014]. Differences in lidar collection specifications, especially lidar pulse density, which varied by roughly a factor of two, might also introduce artifacts if some metrics are particularly sensitive [e.g. Görgens et al., 2015] or are simply hard to reproduce [e.g. metrics based on lidar intensity, Bater et al., 2011]. That said, canopy height and cover metrics used in this study are likely relatively stable across acquisitions, and the LANDSAT data used in our model were collected during the sampling period, with a view to capturing species’ niche axes such as vegetation phenology, habitat type and condition [Leitão and Santos, 2019]. An open question for future studies is whether it is better to include only the individual satellite spectral bands and let the DNN combine the bands, rather than also including known band combinations like NDVI.

#### Choice of JSDM software and interpretation

Our choice of sjSDM over other JSDM software packages was largely dictated by sjSDM’s much faster runtimes while exhibiting predictive performance levels that match other packages [Pichler and Hartig, 2021]. sjSDM also uniquely provides the option to use a combination of regularization and a deep neural network for model fitting, which is appropriate for situations with large numbers of environmental covariates, such as our use of remote-sensing layers, and where the focus is on the predictive power of a model. To compare the effect of using a DNN, we reran the sjSDM model with the same setup but linear in the environmental part. The explanatory AUC of the linear model is higher than in the DNN model, but the predictive power is lower, showing more overfitting with the linear model (Figure 11S). A DNN fitting procedure thus appears to be useful for disentangling complex relationships between remote-sensing-derived environmental covariates and community data.

Going forward, new JSDM software packages are being published that can exploit sample replication to account for false negatives and false positives [Diana et al., 2022, Tobler et al., 2019, Doser et al., 2022]. Over time, as such capabilities are combined with increased efficiency, the result should be more reliable predictions.

Finally, joint species distribution models are distinguished by estimating not only species responses to environmental covariates (as in all species distribution models) but also by estimating correlations between all species pairs while accounting for environmental responses. These residual species associations can be interpreted as the effect of unmeasured environmental covariates and/or the effect of biotic interactions, such as competition or facilitation [Ovaskainen et al., 2017, Pollock et al., 2014, Warton et al., 2015]. It has proven difficult to distinguish between the two in practice [Dormann et al., 2018, König et al., 2021, Poggiato et al., 2021, Zurell et al., 2018, Hartig et al., 2023], and in this study, we are agnostic as to the interpretation of residual species correlations.

## Additional Supplementary Figures and Tables

**Table 1S.**
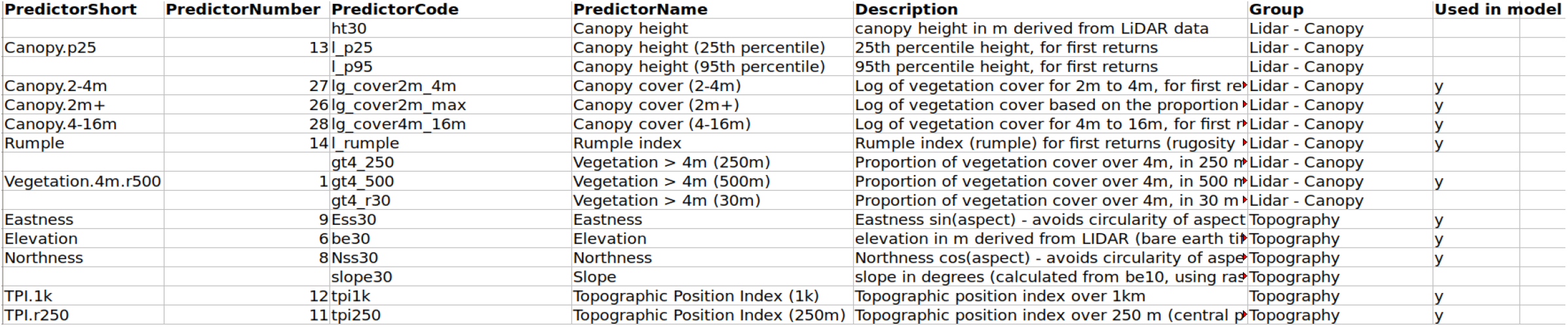
All candidate predictors for jSDM model. Predictors are grouped by origin: Lidar, Landsat, H.J. Andrews Experimental Forest GIS data; 29 predictors were included in the model, chosen by Variance Inflation Factor (VIF) *<* 8, as well as the categorical predictor of inside or outside the boundaries of H.J. Andrews Experimental Forest. Elevation was forced to be included regardless of VIF value. The full table is in https://github.com/chnpenny/HJA_analyses_Kelpie_clean/blob/main/05_supplement/GIS/Table_1S.xlsx

**Figure 3S.**
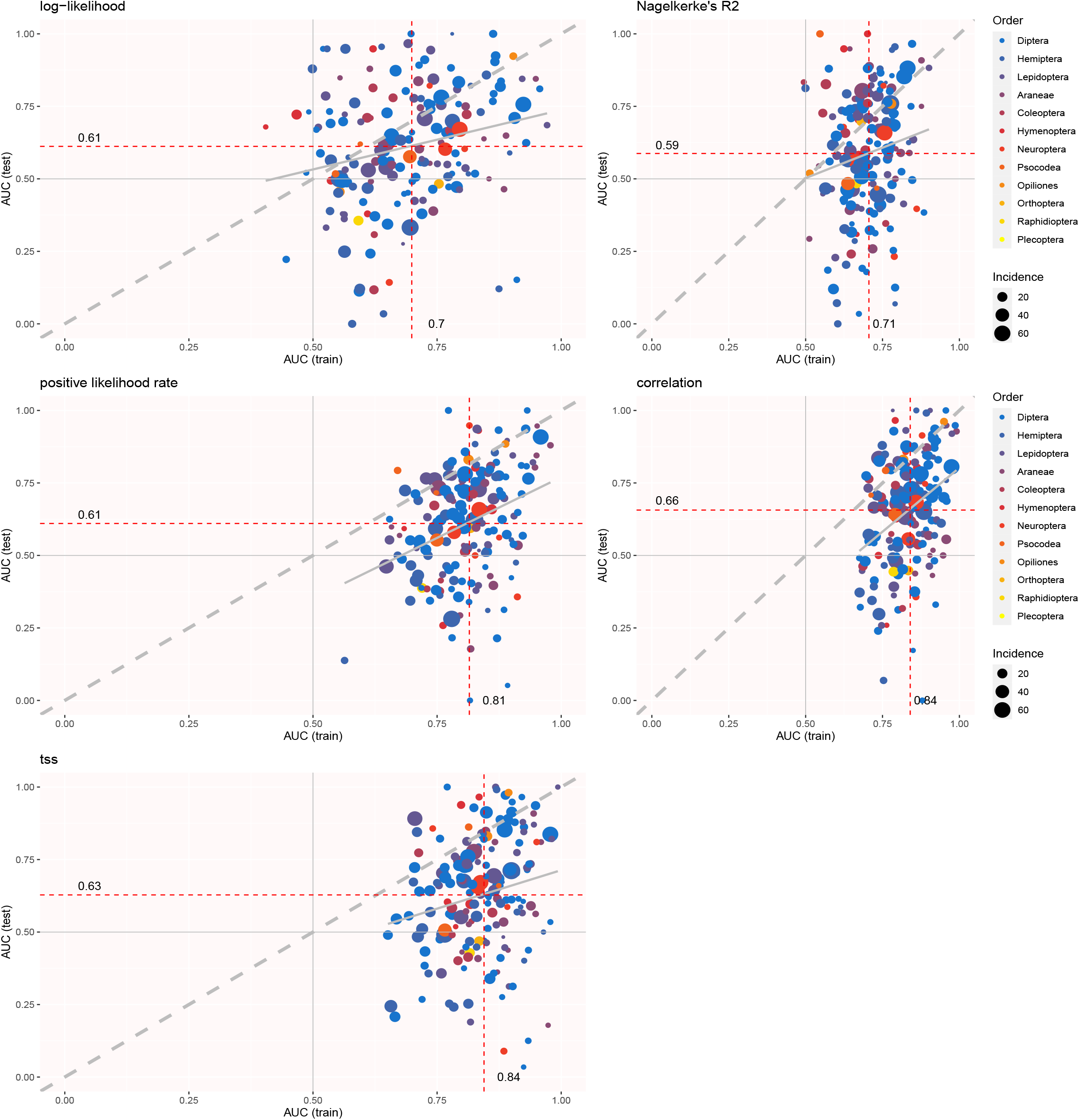
Explanatory AUC vs predictive AUC for best sjSDM models tuned according to log-likelihood, Nagelkerke’s R^2^, positive likelihood rate, correlation and TSS(true skill statistic). Each point is one OTU. Color indicates taxonomic class (order), and point size indicates incidence (number of Malaise traps in which the OTU was detected). The dashed gray line is the 1 : 1 line, and the solid gray line is a fitted linear regression.

**Figure 4S.**
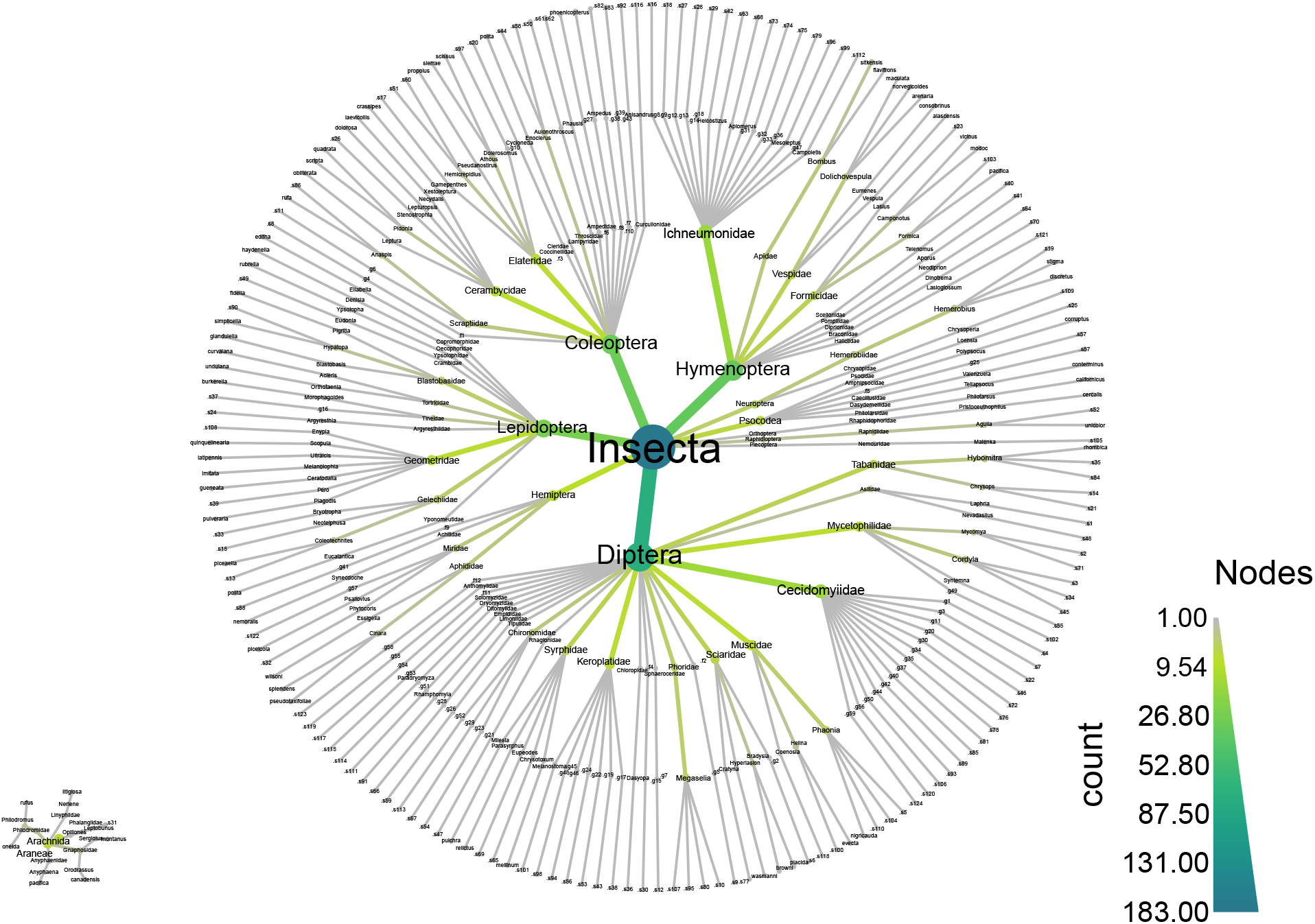
Detailed taxonomic distribution of 190 Operational Taxonomic Units (OTUs) over two heat trees, the Insecta and the Arachnida. Node size and color are scaled to the number of OTUs in that node. Missing taxonomic information of species are indicated by the combination of a point, f, g or s, representing family, genus or species, respectively, and a number, e.g. ‘.f15’.

**Figure 5S.**
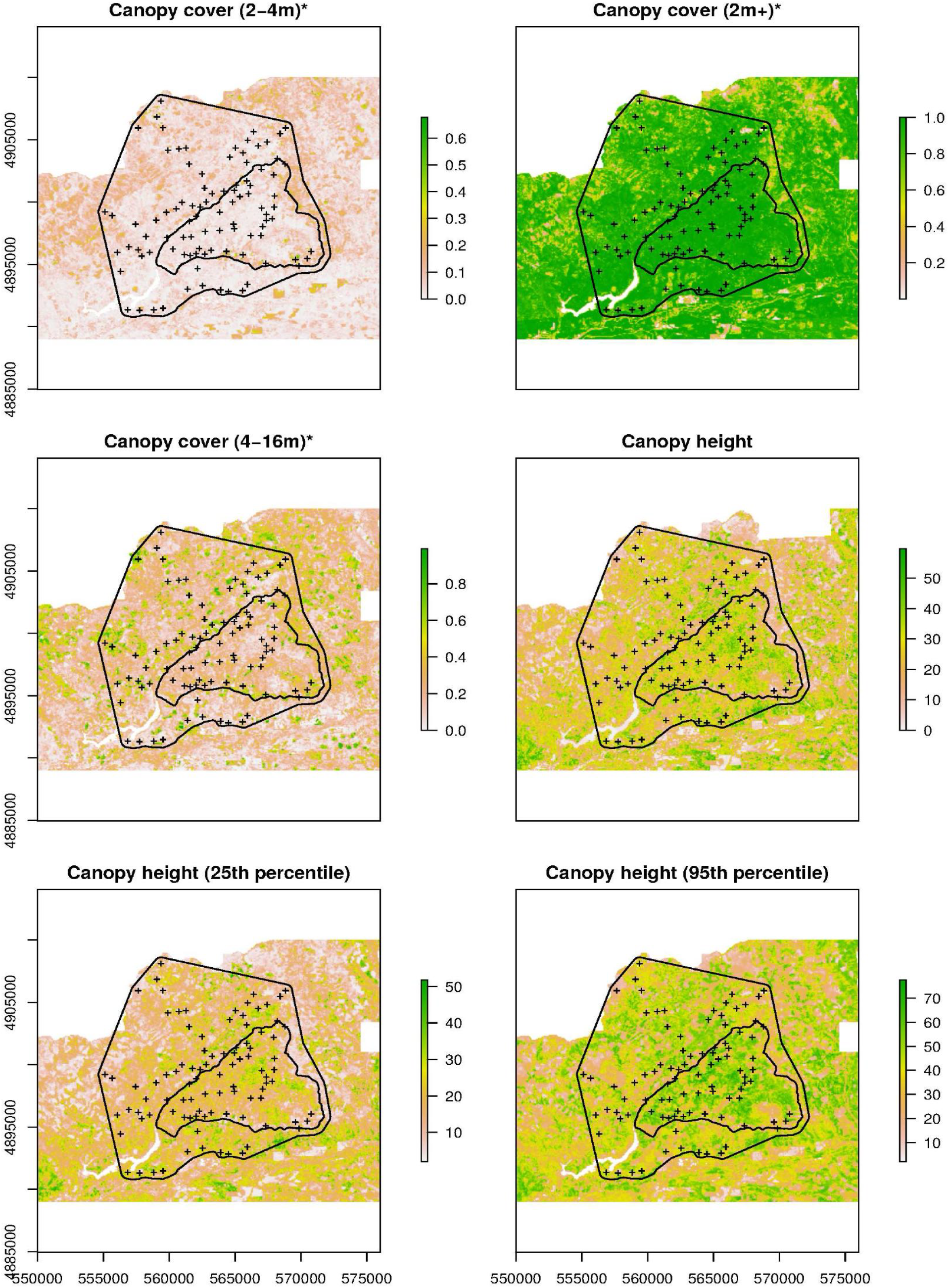
All candidate covariates. Sample locations are marked by the plus sign, inner black outline shows H.J. Andrews Experimental Forest boundary and outer black outline shows extent of prediction area, Covariates used in model are marked with an asterisk. See Table S-covariates for covariate descriptions. The full figures are in https://github.com/chnpenny/HJA_analyses_Kelpie_clean/blob/main/05_supplement/Plots/Figure_5S-full.pdf.

**Figure 6S.**
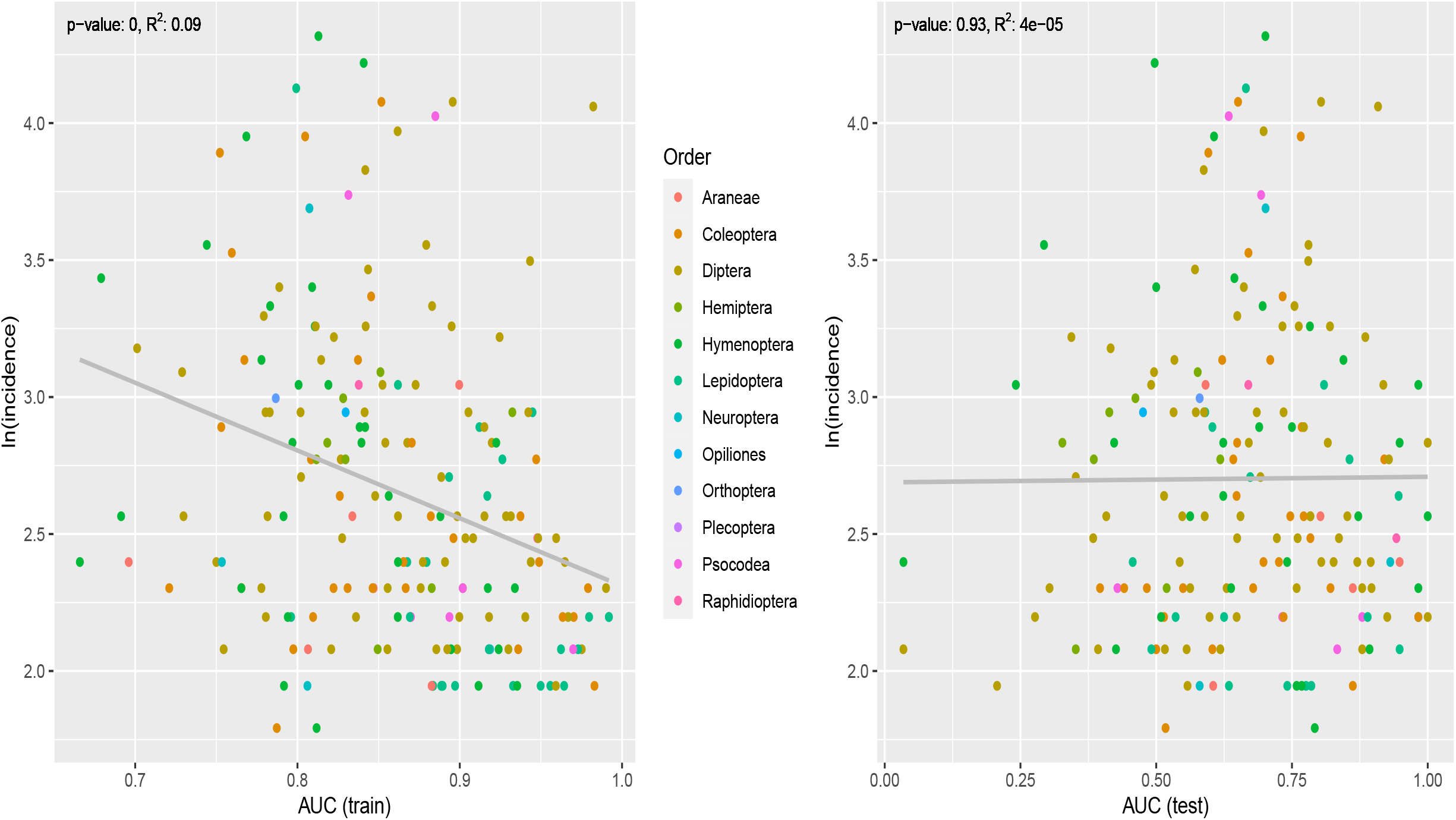
Explanatory (training) and predictive (test) AUCs of all OTUs by incidence. Colors correspond with the order of OTUs. OTUs that are detected less (low incidence) show larger variance in the AUC values. The p-value and *R*^2^ of the linear regressions are shown on the top of the plots. To be noticed, incidences of OTUs are log-transformed.

**Figure 7S.**
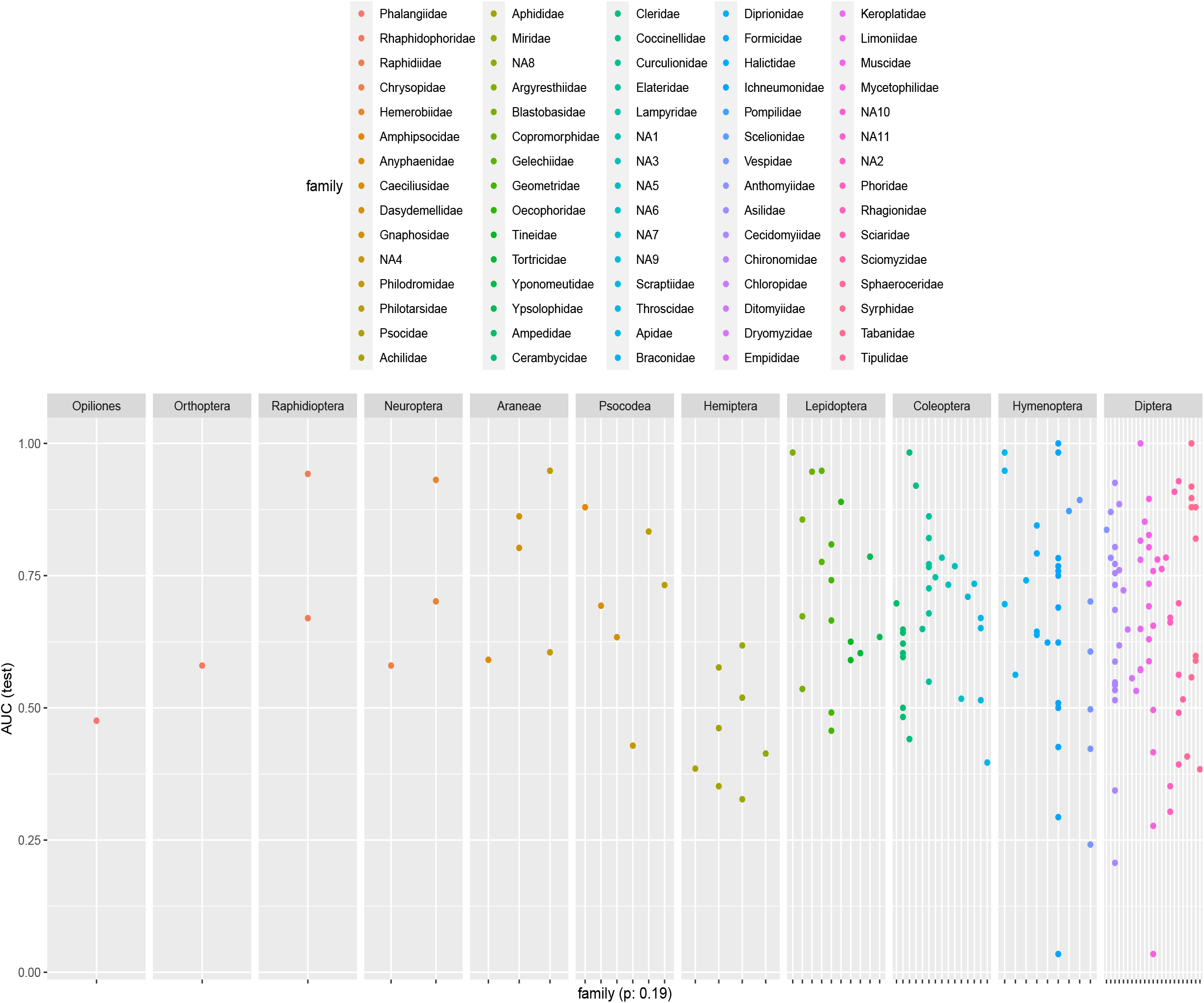
Predictive AUCs of all OTUs by taxonomic family. Colors correspond with the family information and they are arranged according to the order information. A linear regression shows that there is no significant effect of family on the predictive AUCs (p-value 0.19 for this regression).

**Figure 8S.**
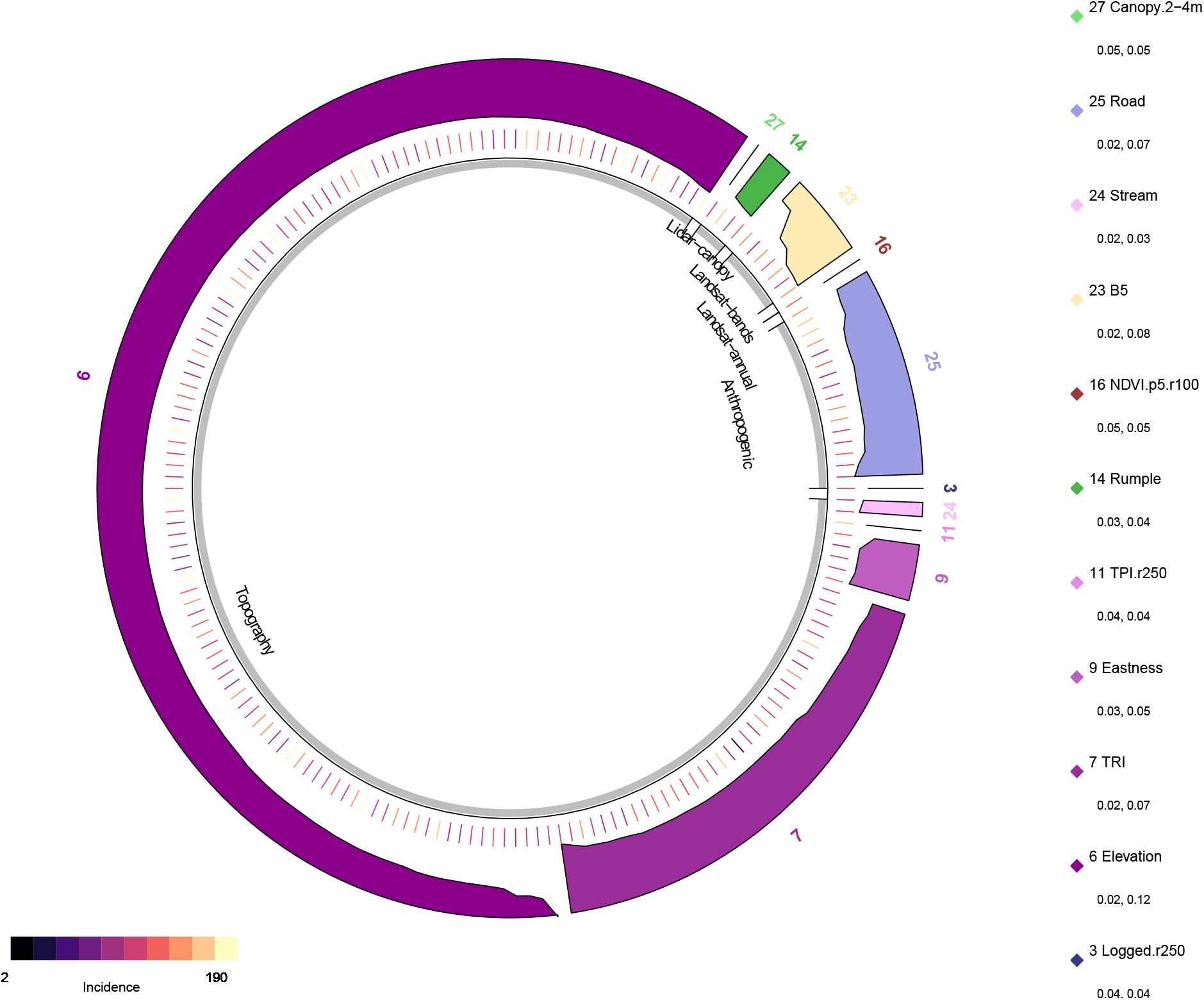
The most important environmental covariate with regard to interaction effects for each OTU, excluding spatial location variables. Each tick mark on the middle ring represents an OTU, coloured by its incidence (see legend lower left), with the outer colour bands indicating its most important individual covariate from the point of view of interaction strength. The effect of environmental covariates on species (i.e. OTU) distributions is comprised of its individual effect and its effect through interacting with other covariates (detail in section, Variable importance with explainable AI (xAI)). Gray bands in the inner ring indicate covariate groupings (Table 1S). Elevation (variable 6) and TRI (variable 7) are the most important variables for the most OTUs. The heights of the colour bands are scaled to the Friedman’s H statistic for overall interaction strength for that OTU. The ranges of overall interaction strength for each environmental variable are shown in the legend on the right.

**Figure 9S.**
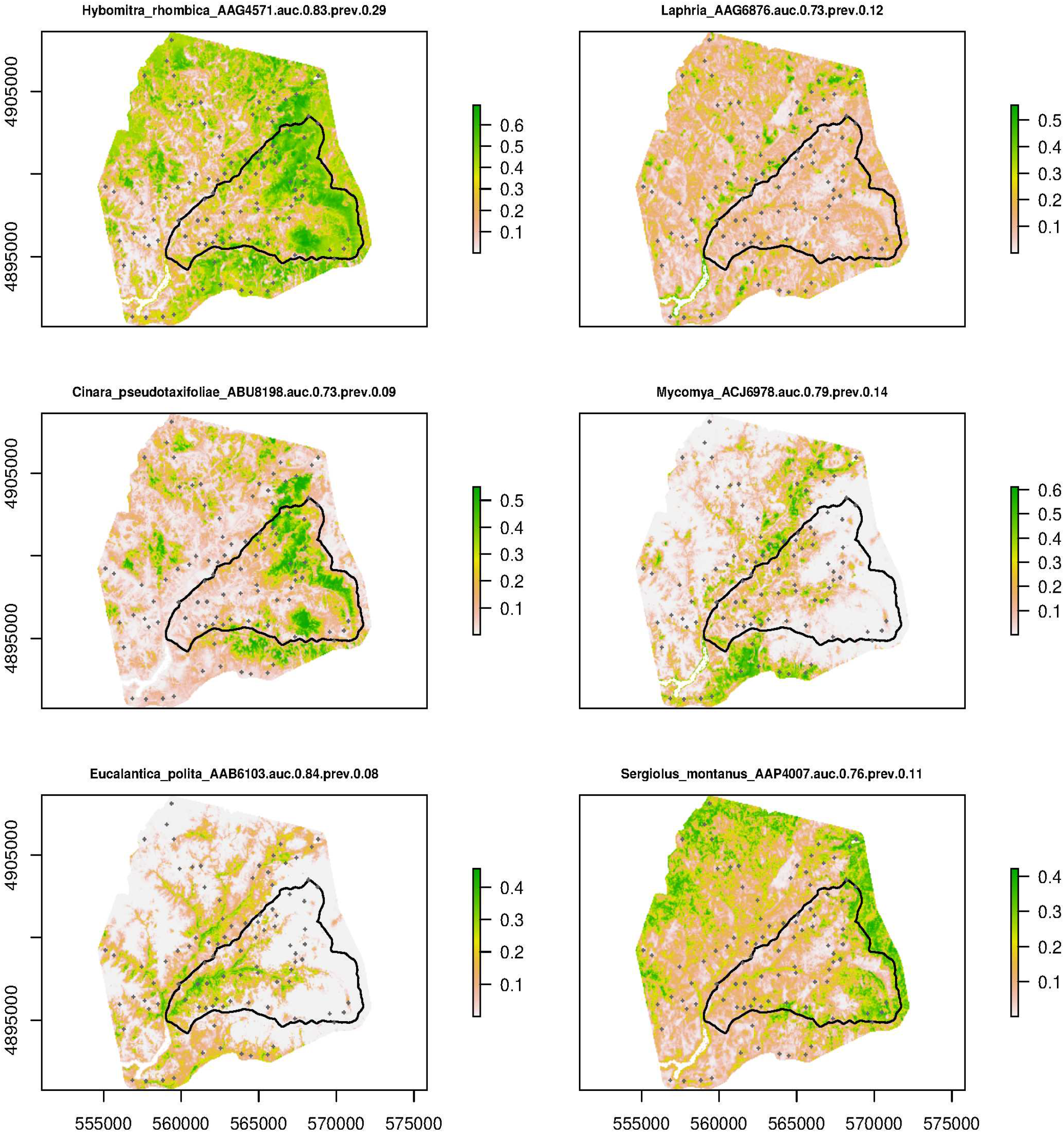
Individual, interpolated species distributions. The full figure is in https://github.com/chnpenny/HJA_analyses_Kelpie_clean/blob/main/05_supplement/Plots/Figure_9S-full.pdf

**Figure 10S.**
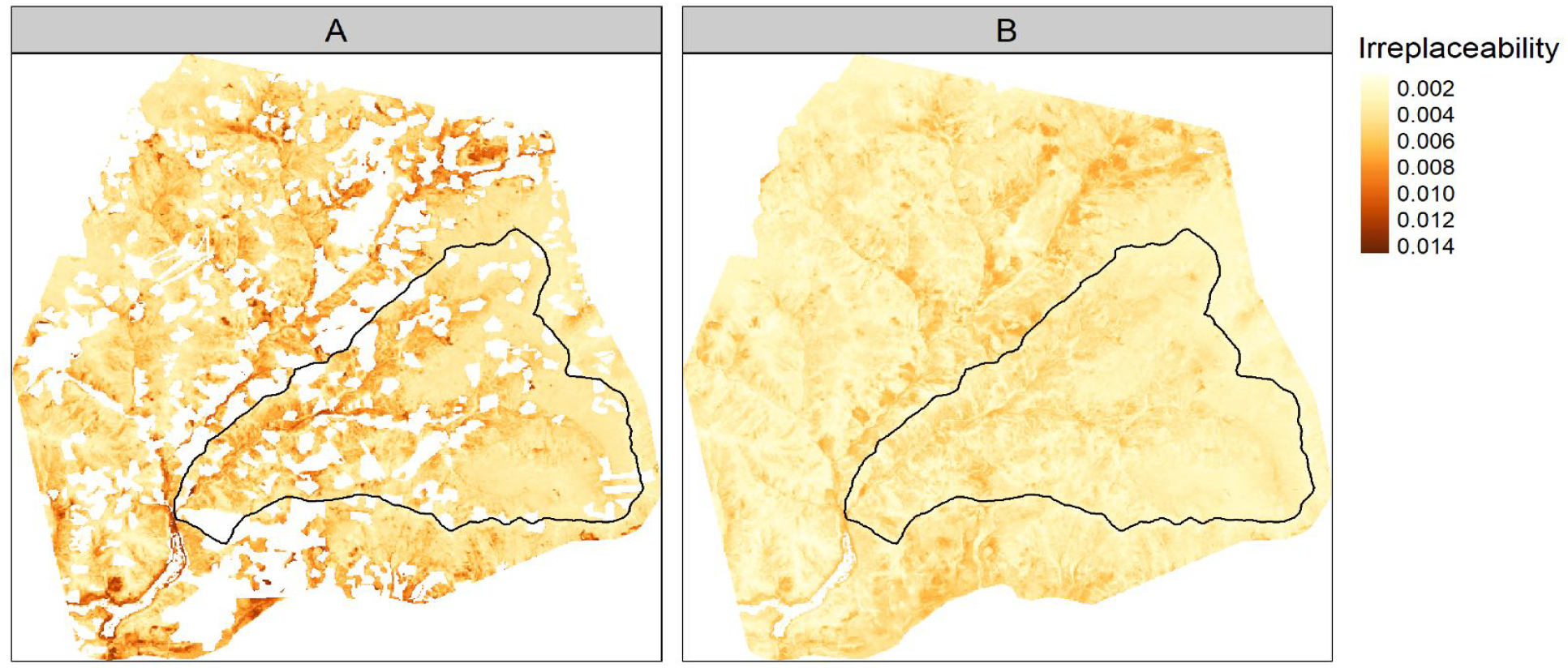
Site-irreplaceability values plotted across the study area, showing HJA Experimental Forest boundaries (black line). A. With plantations masked out. B. With plantations present. Note the higher irreplaceability values in the unmasked part of the landscape (mostly along stream courses), which is because the species that are mainly restricted to plantations are rarer across our study area than those in old growth forests.

**Figure 11S.**
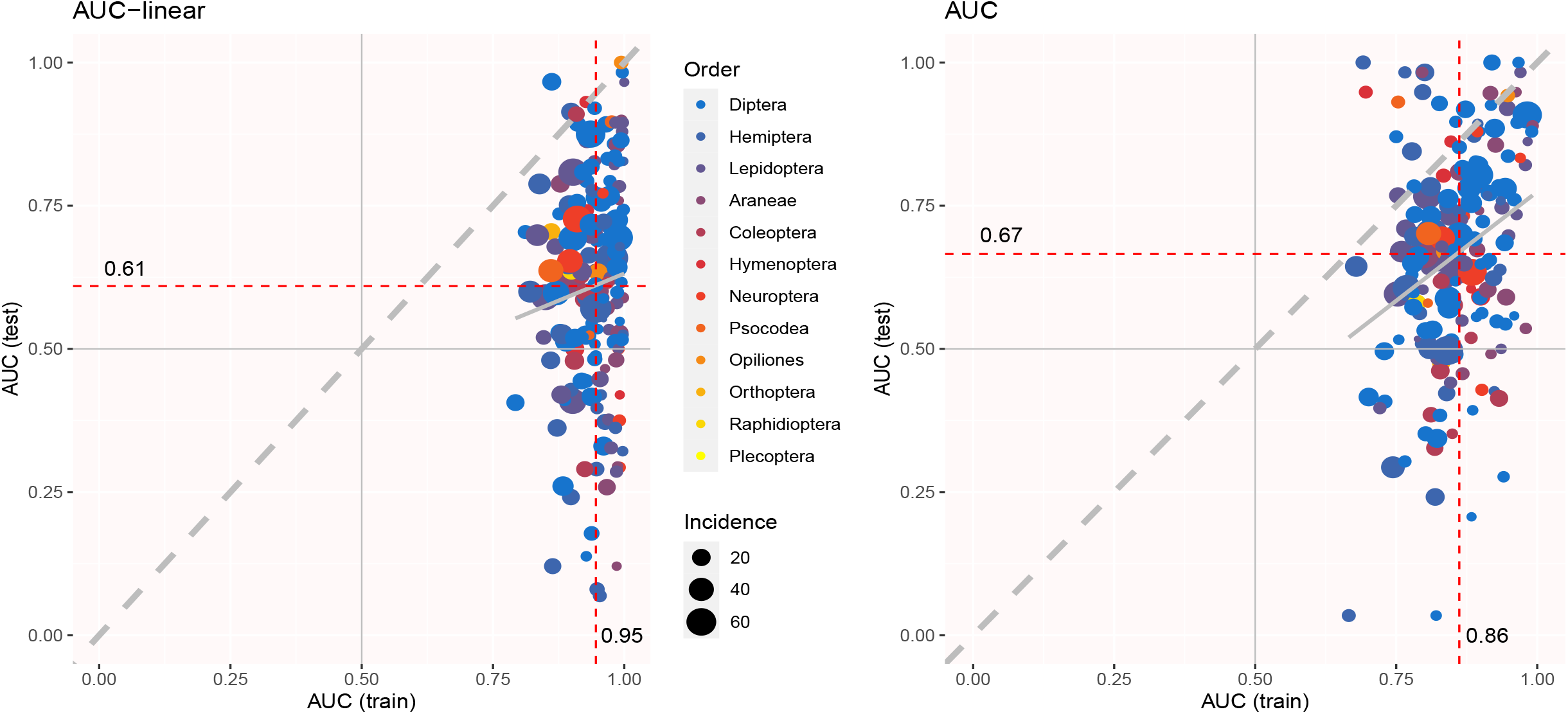
Explanatory and predictive AUCs of the tuned sjSDM model applying linear fitting on the environmental part (left panel) to the same model applying DNN fitting (right panel). The explanatory power (x axis, AUC (train)) is higher but the predictive power (y axis, AUC (test)) is lower in the linear model, relative to the DNN model.

**Figure 12S.**
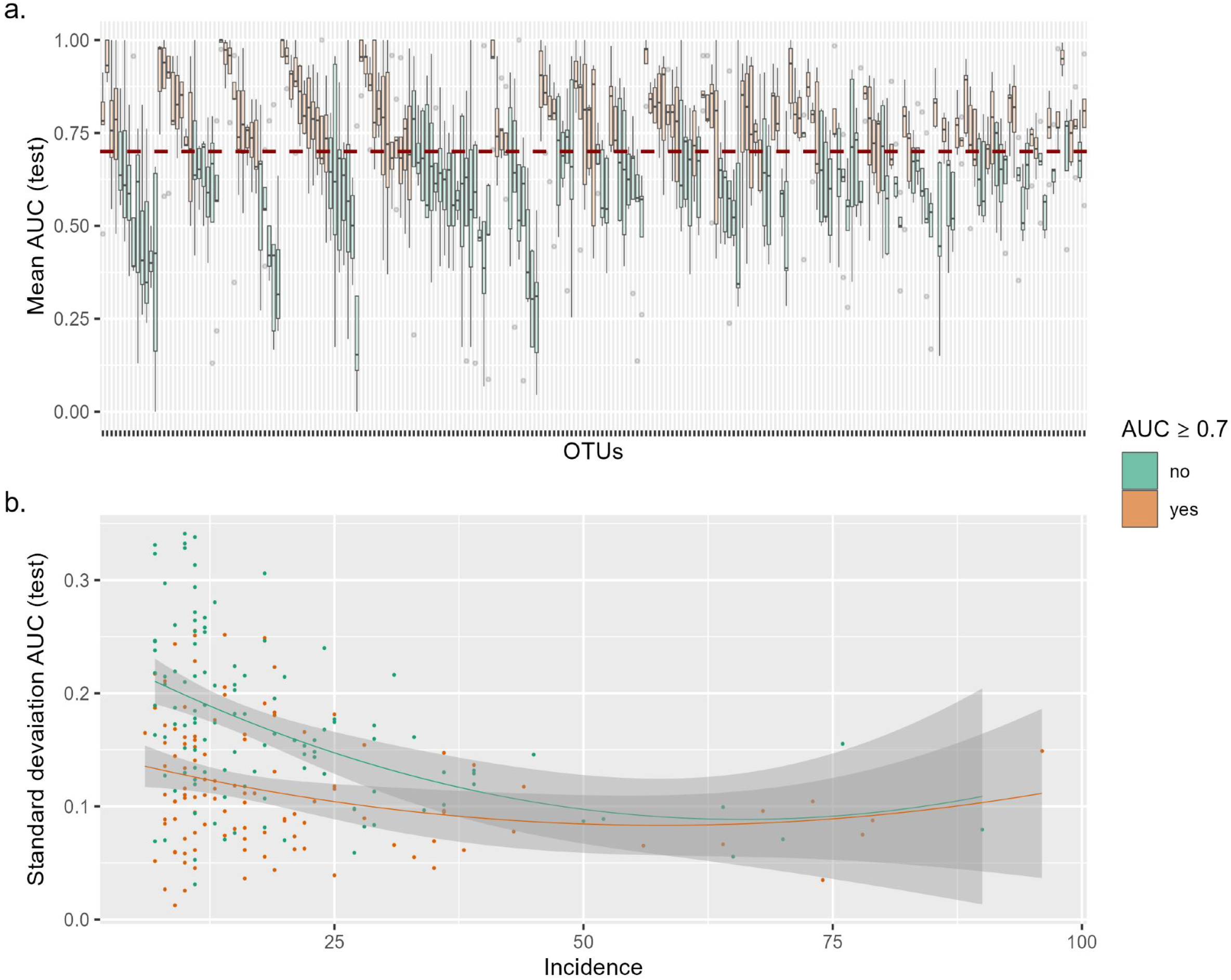
Variability in AUC scores for all OTUs as evaluated with 5-fold cross validation. Variability in AUC is only weakly higher for lower incidence OTUs, and mean AUC does not increase with higher incidence. a) OTUs (boxes) in orange have *AUC*_*mean*_ ≥ 0.70, and those in green have *AUC*_*mean*_ *<* 0.70. OTUs are ordered by increasing incidence, from occurrence at 6 sample points (far left) to occurrence at 96 sample points (far right). The dashed red line is at *AUC* = 0.70, which is the threshold value for including OTUs in further analysis. b) Standard deviation of AUC as a function of incidence. Regression lines shown from a polynomial linear model on OTUs with *AUC*_*mean*_ ≥ 0.70 (*R*^2^ = 0.05, *p* = 0.029, *df* = 2, 109) and with *AUC*_*mean*_ *<* 0.70 (*R*^2^ = 0.19, *p <* 0.001, *df* = 2, 110).

**Figure 13S.**
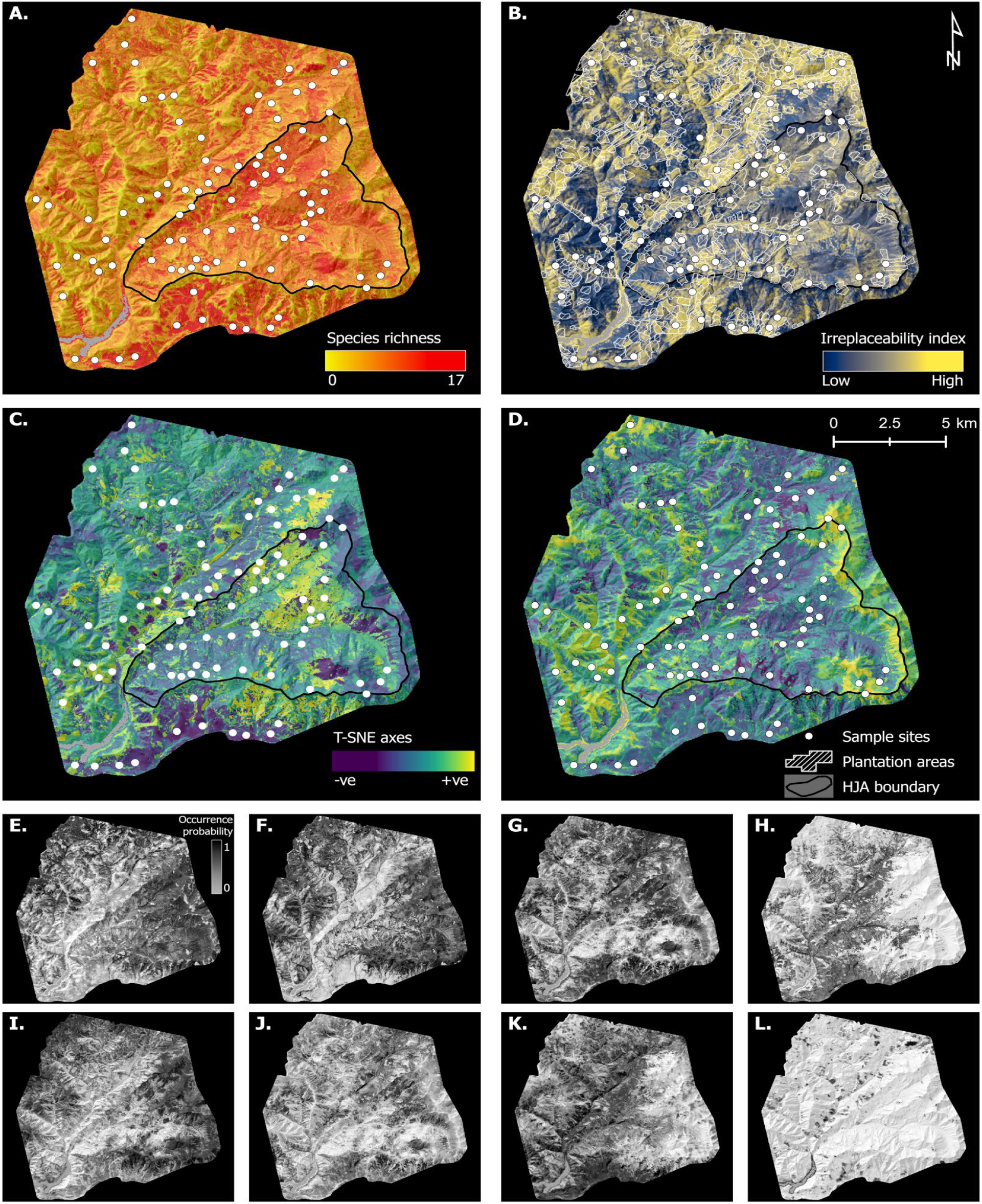
Alternative version of Figure 2 (main text) using OTUs with *AUC*_*mean*_ ≥ 0.70 from 5-fold CV on full data set (see methods in Tuning and Testing, above). JSDM-interpolated spatial variation in species richness, irreplaceability, and composition, plus examples of individual species distributions. A. Species richness. B. Site beta irreplaceability, showing areas of forest plantation. C-D. T-SNE axes 1 and 2. White circles indicate sampling points, white polygons indicate plantation areas (i.e. a record of logging in the last 100 years), and the black-line-bordered triangular area delimits the H.J. Andrews Experimental Forest (HJA, see Figure 1, main text). E-L. Selected individual species distributions, with BOLD ID, predictive AUC, and prevalence. E. Rhagionidae gen. sp. (BOLD: ACX1094, AUC: 0.95, Prev: 0.64). F. *Plagodis pulveraria* (BOLD: AAA6013, AUC: 0.72, Prev: 0.23). G. *Phaonia* sp.(BOLD: ACI3443, AUC: 0.80, Prev: 0.65). H. *Orthotaenia undulana* (BOLD: AAB4022, AUC: 0.95, Prev: 0.06). I. *Helina evecta* (BOLD: AAC2498, AUC: 0.76, Prev: 0.16). J. *Diptera* sp. (BOLD: AAZ4857, AUC: 0.75, Prev: 0.16). K. *Blastobasis glandulella* (BOLD: AAG8588, AUC: 0.91, Prev: 0.18). L. *Dasyopa* sp. (BOLD: ADI1308, AUC: 0.82, Prev: 0.12)

## Notes

https://github.com/chnpenny/HJA_analyses_Kelpie_clean/releases/tag/v1.1.0

https://zenodo.org/record/8303158

### Competing Interest Statement

DWY is a co-founder of NatureMetrics (www.naturemetrics.com), which provides commercial metabarcoding services.

### Summary of Updates

Author-accepted manuscript (Philosophical Transactions of the Royal Society B)

